# LoxCode in vivo barcoding resolves epiblast clonal fate to fetal organs

**DOI:** 10.1101/2023.01.02.522501

**Authors:** Tom S. Weber, Christine Biben, Denise C. Miles, Stefan Glaser, Sara Tomei, Michael Lin, Andrew Kueh, Martin Pal, Stephen Zhang, Patrick P. L. Tam, Samir Taoudi, Shalin H. Naik

**Affiliations:** Immunology Division, WEHI, Parkville, Melbourne. Australia. 3052; Department of Medical Biology, University of Melbourne, Melbourne, Australia. 3052; Epigenetics and Development Division, WEHI, Parkville, Melbourne. Australia. 3052; Boehringer Ingelheim RCV GmbH & Co KG, Austria; Blood Cells and Blood Cancer Division, WEHI, Parkville, Melbourne. Australia. 3052; Olivia Newton John Cancer Research Institute, 145 Studley Road, Heidelberg, Victoria 3084, Australia; School of Cancer Medicine, La Trobe University, Bundoora, Victoria 3086, Australia; School of Dentistry and Medical Sciences, Charles Sturt University, Wagga Wagga, New South Wales, Australia; Embryology Research Unit, Children’s Medical Research Institute, University of Sydney. Westmead, NSW 2145, Australia; School of Medical Sciences, Faculty of Medicine and Health, University of Sydney, Sydney NSW 2006, Australia

## Abstract

Much remains to be learned about the clonal fate of mammalian epiblast cells *in vivo*. Here we develop a high diversity, high throughput, Cre recombinase-driven DNA LoxCode barcoding technology for *in vivo* clonal lineage tracing. E5.5 pre-gastrulation embryos were barcoded *in utero* and epiblast clones later assessed for their contribution to a wide range of tissues and cell types in the E12.5 organogenesis-stage embryo. While a few epiblast clones contributed broadly to most tissues and cell types of the three germ layers, many clones were lineage biased towards either blood, ectoderm lineages, mesenchymal tissues or limbs. In addition to lineage bias, most epiblast clones were differentially fated for tissue types or descendants in tissues compartments across the body axes. Using a stochastic agent-based model of embryogenesis and LoxCode barcoding, we inferred and experimentally validated predicted cell fate biases as well as clonal compositions across tissues that are consistent with events of lineage segregation and shared trajectory of lineage differentiation. Our study has demonstrated the power of LoxCode barcoding in investigating multi-modal clonal fate at high throughput, thus enabling an in-depth interrogation of the clonal contribution of E5.5 epiblast cells to fetal tissues, organs and body parts.

## Introduction

Mammalian embryogenesis involves the generation and assembly of different cell and tissue types derived from the epiblast into a body plan that constitutes the blueprint of embryonic development. However, it is unclear whether all epiblast cells are multipotent or whether certain cells are already lineage restricted prior to gastrulation. Answering this question in the mammalian embryo has been hampered by the inability to track cells of the epiblast at single-cell resolution over an extended timeline in vivo. Seminal studies have shed light on the early stages of mouse embryogenesis, including the production of a presumptive fate map for the epiblast embryo^1^, the discovery of how spatial position of cells in the epiblast influences the acquisition of developmental fate^2^, single-cell RNA sequencing and fluorescent in situ hybridization of embryonic cells and tissues at different developmental stages ^3-9^, and the lineage tracing of single cell clones of the epiblast during gastrulation^10,11^. Despite the advances made in these studies of prospective cell fate, including a more recent *in vivo* pedigree analysis^5^, technical challenges remain for implementing clonal lineage analysis through both prospective and/or retrospective tracing of lineage trajectories *in vivo* across gastrulation to organogenesis^6^. Parallel assessment of the development of multiple clones is imperative to answer such questions.

Technological advances in cellular barcoding have empowered the tracing of clonal lineages^12,13^. Hereby, a progenitor population of cells is tagged with unique and heritable barcodes prior to further proliferation and differentiation. Subsequent comparison of the barcode repertoire of multiple clones contributing to cellular progeny allows determination of clonal relationships, e.g., derivation from common versus separate ancestors, estimation of clone sizes, timing of lineage allocation and diversification, and spatial location of the lineage precursor. Unlike viral barcoding methods, that transduce cells *ex vivo* which are then engrafted into a recipient, recent *in vivo* technologies enable barcode generation in cells in their native environment. This feature offers the ability to investigate clonal biology in a *native* setting. These barcoding techniques encompass prospective (multispectral barcoding^14^, *in vivo* viral delivery^15^, transposons^16,17^, CRISPR/Cas9^5,18-22^, Cre/Lox recombination^23,24^) and retrospective (natural barcodes^25^, cell history recorder^26,27^) approaches. Many also allow the simultaneous assessment of the transcriptome and clonal origin of single cells for leveraging the ability to link cell state to cell fate, or to determine shared molecular features among clones^28^.

Previously, we established the foundational design principles for a Cre/Lox-based barcode generation system and mathematically determined that the arrangement of distinct DNA elements flanked by LoxP sites in alternating directions would lead to optimal barcode diversity^29^. Notably, our findings indicated that the number of theoretical barcodes would increase combinatorically with the addition of each barcode element to the cassette. For example, a configuration comprising 14 LoxP sites and 13 code elements could potentially produce a barcode diversity exceeding 30 billion upon partial recombination (*Methods*). Furthermore, our design incorporated a novel aspect: utilizing DNA elements of minimal length to create a compact construct and limit further excisions once the LoxCode cassette reaches a size of three barcode elements. This feature facilitated efficient recombination at low Cre concentrations, generated over 2,000 size-stable barcodes resistant to further Cre excisions and ensured, in contrast to established Cre based barcoding systems^23^.

Here we describe how LoxCode clonal lineage tracing offers several critical attributes that permit resolution of clonal pattern of cell/tissue lineages. Upon LoxCode generation in the E5.5 epiblast and LoxCode extraction from tissue of E12.5 embryo either through PCR or single cell RNA-seq, and applying a suite of novel computational methods, we have established i) the contribution of epiblast cellular descendants to a full range of embryonic tissues, ii) the inter-relationships of fetal tissues and organs, and iii) a better understanding of cell types derived from individual epiblast cells, which informed iv) cell type relationships at a clonal level. In parallel to the present study, we have applied LoxCode clonal lineage tracing to investigate lineage relationships during yolk sac haematopoiesis^30^.

## RESULTS

### LoxCode design and experimental setup

Based on optimal design principles for Cre/LoxP cellular barcoding^29^, the LoxCode was constructed through serial integration of LoxP/Barcode paired inserts to generate a final cassette of 14 LoxP sites of alternating orientation flanking 13 short barcode elements of 8 bp and 14 bp (Fig. 1a, *Methods*). Upon Cre exposure, random recombination between the LoxP sites led to inversions and excisions, resulting in DNA sequence reshuffling with a theoretical DNA diversity of over 30 billion barcodes (Methods and Table 1). The LoxCode construct was introduced into the *Rosa26* safe harbor locus in C57BL/6 mice to enable *in vivo* lineage tracing though both PCR-based detection in genomic DNA as well as read-out of LoxCodes transcripts driven by the endogenous *Rosa26* promoter (Supplementary Fig. 1).

**Figure 1.**
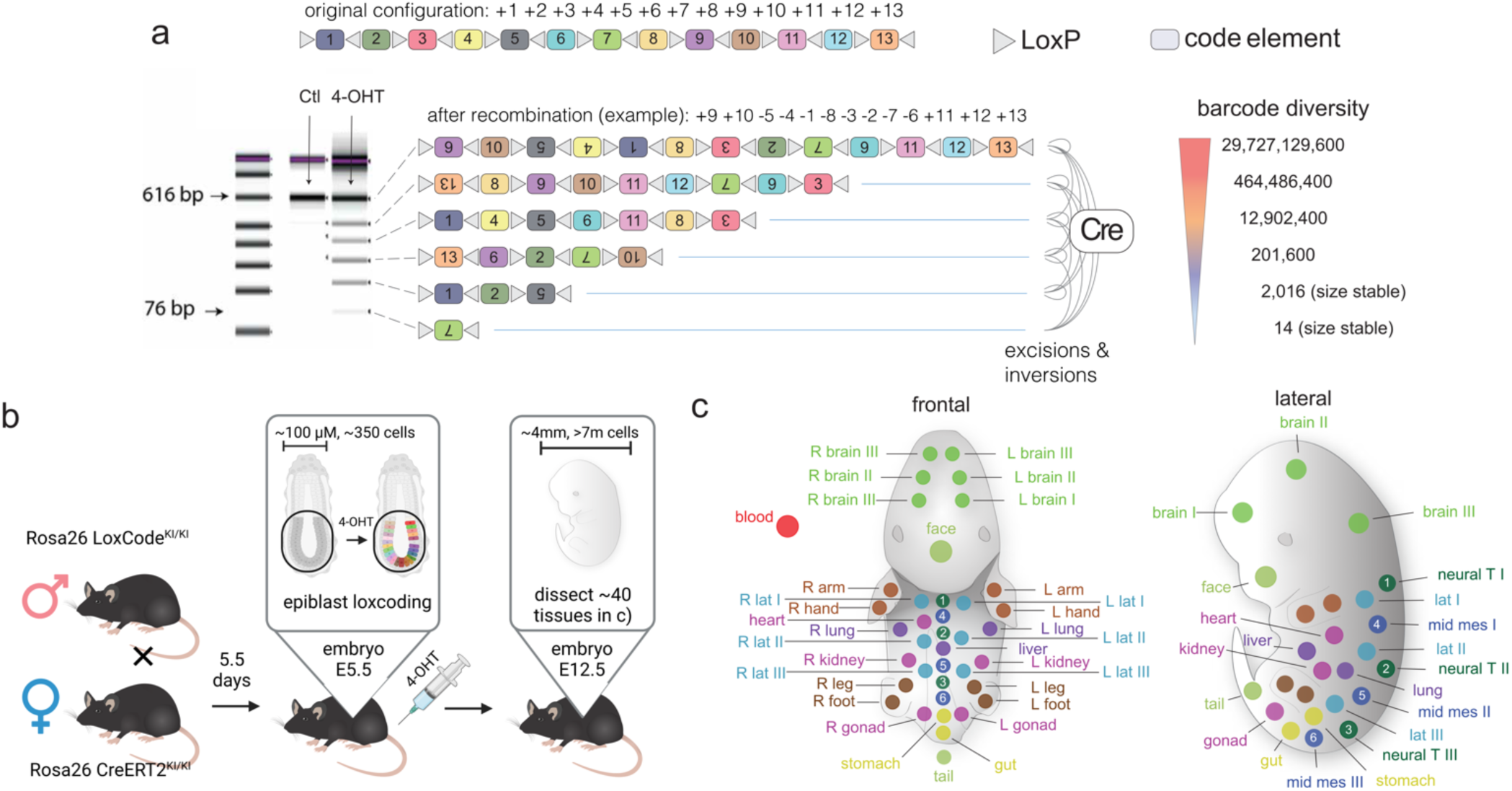
LoxCode design and experimental setup. **a)** The LoxCode cassette is composed of 14 LoxP sites in alternating orientation flanking 13 small (8-14 nucleotides) barcode elements indexed from 1 to 13. Cre exposure leads to random excisions and inversions, and different sized bands can be analysed via gel electrophoresis. Hypothetical LoxCodes of different sizes with inverted barcode elements depicted upside down. To annotate barcodes, elements are listed in order, with a plus or minus sign indicating if the orientation matches or is inverted compared to the original sequence. With this notation, the original cassette is written as ’+1+2+3+4+5+6+7+8+9+10+11+12+13’, and the first recombined example in panel a) is annotated as ’+9+10-5-4-1-8-3-2-7-6+11+12+13’ where inverted LoxCode elements are denoted by a minus (-) symbol. Computed theoretical barcode diversities for each barcode size (containing 13, 9, 7, 5, 3, 1 elements) are provided (right column). Experimental design: **b)** Barcoding of all cells (epiblast cells are highlighted in colour) in E5.5 pre-gastrulation embryos was activated by administration of 4-hydroxytamoxifen (4-OHT) to the pregnant mice, followed by dissection of E12.5 embryos into **c)** 40 tissues and organs for barcode retrieval and single-cell RNA sequencing analysis.

**Table 1.**
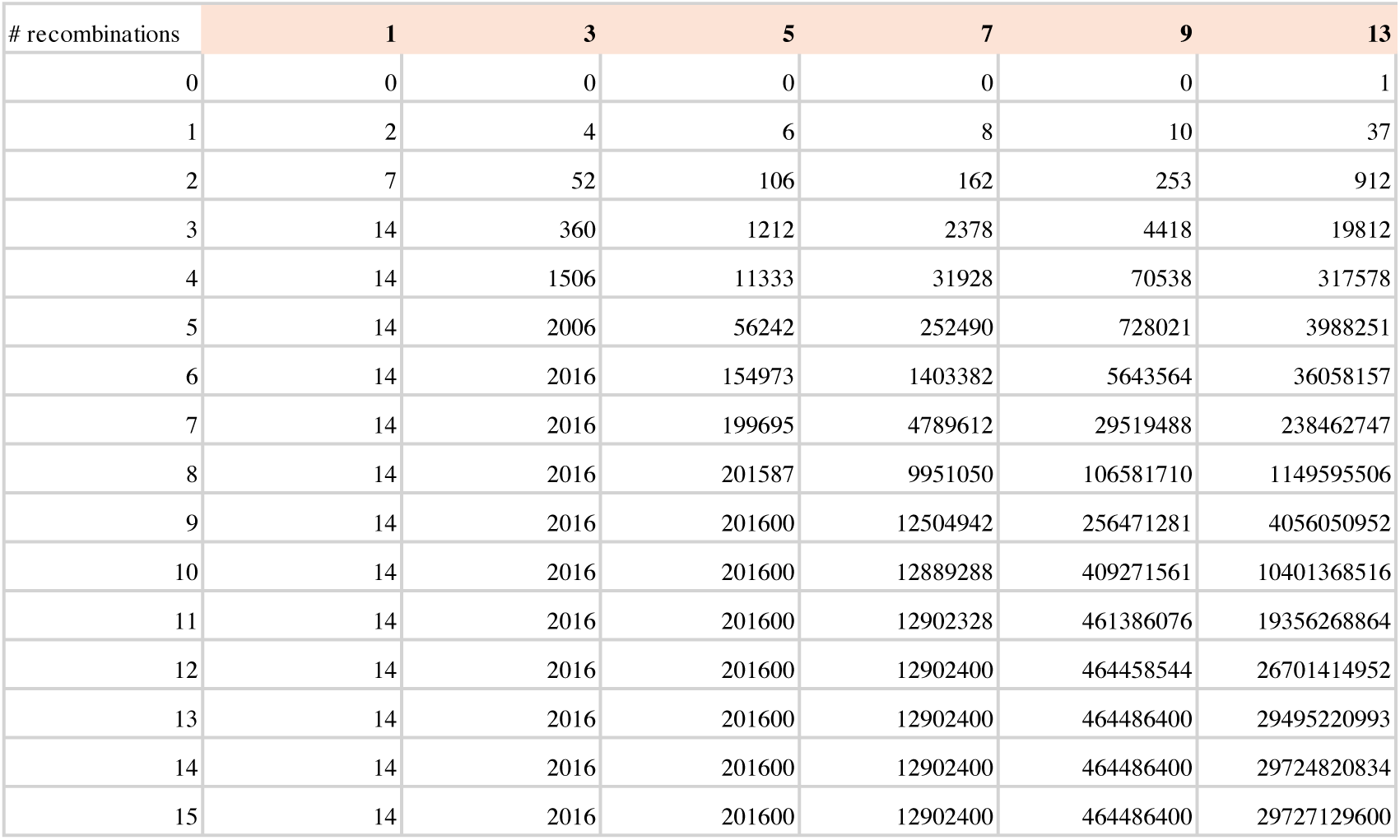
Theoretical barcode diversity for barcodes with 1 to 13 elements (columns) per number of recombination events (rows).

| # recombinations | 1 | 3 | 5 | 7 | 9 | 13 |
| --- | --- | --- | --- | --- | --- | --- |
| 0 | 0 | 0 | 0 | 0 | 0 | 1 |
| 1 | 2 | 4 | 6 | 8 | 10 | 37 |
| 2 | 7 | 52 | 106 | 162 | 253 | 912 |
| 3 | 14 | 360 | 1212 | 2378 | 4418 | 19812 |
| 4 | 14 | 1506 | 11333 | 31928 | 70538 | 317578 |
| 5 | 14 | 2006 | 56242 | 252490 | 728021 | 3988251 |
| 6 | 14 | 2016 | 154973 | 1403382 | 5643564 | 36058157 |
| 7 | 14 | 2016 | 199695 | 4789612 | 29519488 | 238462747 |
| 8 | 14 | 2016 | 201587 | 9951050 | 106581710 | 1149595506 |
| 9 | 14 | 2016 | 201600 | 12504942 | 256471281 | 4056050952 |
| 10 | 14 | 2016 | 201600 | 12889288 | 409271561 | 10401368516 |
| 11 | 14 | 2016 | 201600 | 12902328 | 461386076 | 19356268864 |
| 12 | 14 | 2016 | 201600 | 12902400 | 464458544 | 26701414952 |
| 13 | 14 | 2016 | 201600 | 12902400 | 464486400 | 29495220993 |
| 14 | 14 | 2016 | 201600 | 12902400 | 464486400 | 29724820834 |
| 15 | 14 | 2016 | 201600 | 12902400 | 464486400 | 29727129600 |

To study the clonal fate of E5.5 embryonic cells *in vivo*, LoxCode homozygous females were crossed to *Rosa26*-Cre-ER^T2^ homozygous males to produce heterozygous LoxCode/*Rosa26*-Cre-ER^T2^ embryos (Fig. 1b). LoxCode generation was induced at E5.5 via intravenous injection (IV) of low doses (50-100 µg) of 4-Hydroxytamoxifen (4-OHT) to both minimise developmental toxicity and to allow partial LoxCode recombination, with the latter attribute enhancing barcode diversity and complexity. Barcode induction was expeditious, with the majority of recombination generated by 3 hrs, and stabilised by 24 hrs (Supplementary Fig. 2 a-b). To assess the clonal fate of E5.5 embryonic cells, E12.5 embryos were dissected into 40 tissues and organs (Fig. 1c) which were either: i) lysed, prior to barcode PCR and sequencing (Fig. 2, 3); or ii) dissociated into single cell suspensions for concurrent scRNA-seq and barcode detection (Fig. 5).

**Figure 2.**
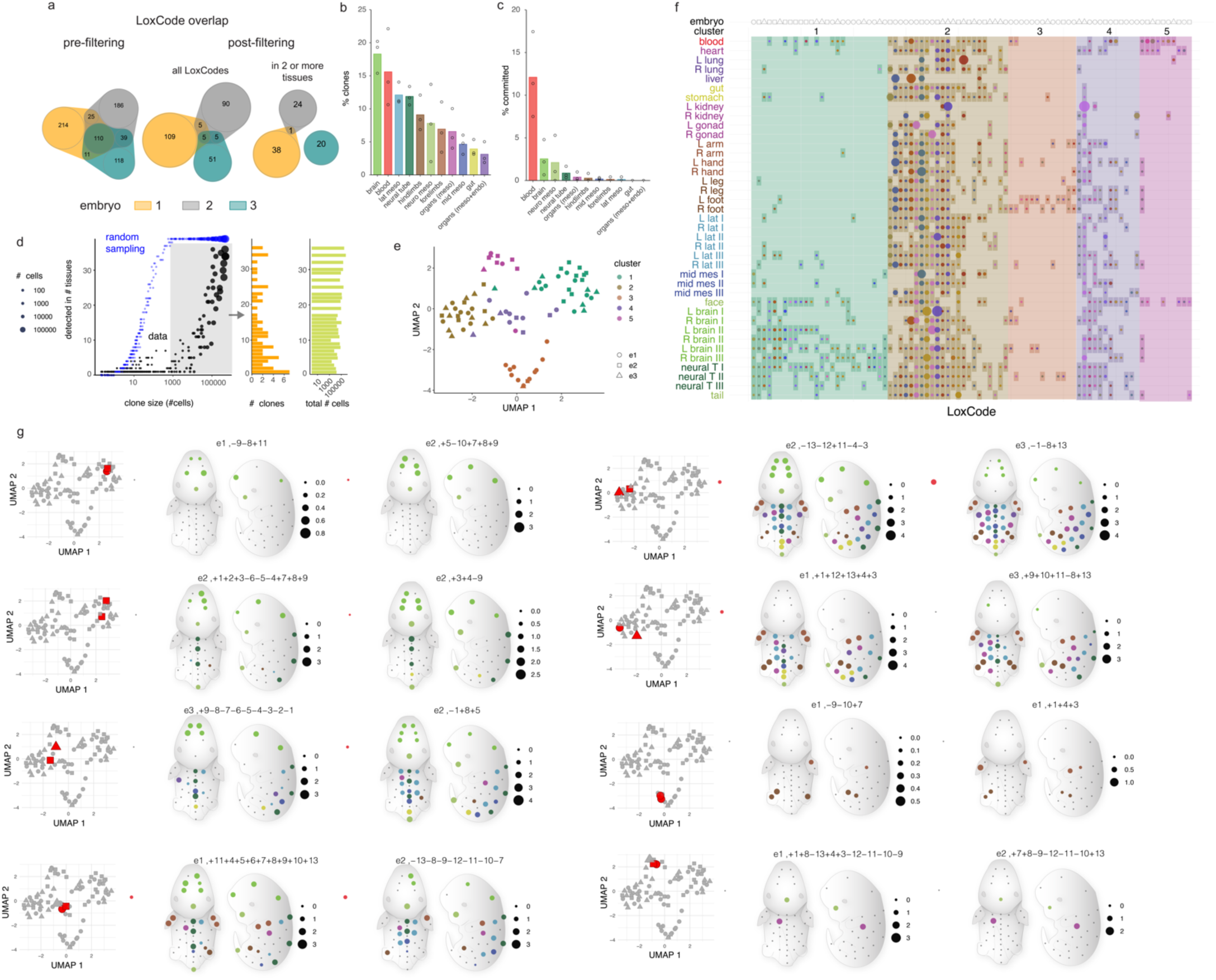
Patterns of epiblast clonal fate to organs and tissues. **a)** LoxCodes were excluded based on empirical frequency (see Methods) resulting in few overlapping barcodes between biological replicates. **b)** Proportion of clones contributing to indicated tissue types. **c)** Proportion of clones uniquely detected in the indicated tissues. d) Comparison of the clone size and number of tissues each LoxCode was detected in, versus a simulation of random sampling of cells from all tissues of the corresponding clone sizes. Histograms on the right show the number of clones and average size for clones with more than 1000 cells. **e)** UMAP projection of clonal outcomes with 5 clusters of clonal output. **f)** Bubble plot/binary heatmap of clonal contribution to embryonic tissues. LoxCodes (columns) are ordered by 1D UMAP according to patterns of their clonal output to the indicated tissues (rows) from 3 embryos. Darkened rectangles indicate all detected clones (binary visualisation), and circles inside are proportional to clonal output to the indicated tissues (quantitative visualisation). Coloured backgrounds (six) correspond to the cluster annotations from e)**. g)** Examples of clonal fate in a pseudo-anatomical map, where circles are scaled proportional to the logarithm of frequency of the indicated LoxCode and preceded by embryo of origin e1, e2, e3, respectively).

While barcoding of the E5.5 embryo using *Rosa26*-Cre-ER^T2^ would not be limited to the epiblast cells, established knowledge of lineage development during embryogenesis indicates that the extraembryonic ectoderm has no or little contribution to the germ layer tissues while a sub-population of visceral endoderm may contribute to the endoderm of the gastrulating embryo^31,32^. Accordingly, we directed our analysis to clones in the late organogenesis stage of the embryo, on the assumption that most clonal descendants detected in the embryo by this stage are derived from epiblast cells.

### Epiblast clonal fate heterogeneity

From three embryos, we recovered a total of 703 barcodes present in both PCR replicates of each tissue (∼200-300 barcodes per embryo similar to the number of epiblast cells at the time of barcoding) of the expected range of barcode sizes. The inferred number of recombination events (excisions and/or inversions) for each barcode ranged from one up to eight (Supplementary Fig. 2 c-d). After removal of frequent barcodes that may arise from more likely recombination events (‘repeat use’ or ‘barcode collisions’) using an empirically derived exclusion list (see Methods) ∼30% of barcodes passed filtering, with an overlap of 10 barcodes between any two embryos, and only 1% (5 out of 265) present in all three embryos (Fig. 2a). Furthermore, clones contributing to at least two tissues represented one third of those barcodes, with very little overlap between embryos. This indicated efficient enrichment of *bona fide* single-use barcodes (Fig. 2a).

In a multipotent model where every epiblast clone contributes broadly to a wide range of cell/tissue types, albeit with varying clone size, one would expect every barcode to be detected in most 40 different tissues or body parts. While we detected a few multi-lineage clones in each embryo (∼5% of all clones), this was not the general rule: allocation of E5.5 epiblast cells ranged from only ∼3.5% clones towards the gut up to a maximum of ∼20% of all clones towards the brain (Fig. 2b). A minority of clones were restricted to a single tissue type which ranged from 1-3% of all clones for most tissues. Strikingly, a significant fraction (12%) of LoxCodes were present exclusively in the circulating blood cells, which at E12.5 is predominantly extra-embryonic yolk sac-derived erythroid cells^33^, indicating particularly early fate restriction (Fig. 2c). This is in line with previous studies^34,35^.

Overall, the majority of LoxCodes of varying clone sizes (spanning three orders of magnitude) were unequally detected in a restricted number of tissues (Fig. 2d), inconsistent with a null-model in which single-cell epiblast fate contributes homogeneously to all tissues. This observation was further corroborated by performing simulations where cell numbers equivalent to the empirically derived clone sizes were drawn randomly from the distribution of cells across all 40 tissues, showing a stark difference between expected clone sizes under the null model and the empirical data (Fig. 2d, Kolmogorov–Smirnov, p-value<0.001).

Next, we explored whether, from the large number of theoretical clonal fates, a limited set of reproducible fate patterns were realised *in vivo*. For this, epiblast clones detected in at least two tissue samples were classified by Leiden clustering^36^ into five main clusters of fate according to their contribution to 11 major tissue types (Fig. 2b) and projected onto 2 dimensions by using the Uniform Manifold Approximation and Projection (UMAP^37,38^) algorithm (Fig. 2e) and summarized in a bubble plot (Fig. 2f). We observed neural tissue-biased (cluster 1), multi-potent (cluster 2), limb-biased (cluster 3), and blood-biased (cluster 5) clones. Cluster 4 clones contributed to many tissues, but less to splanchnopleure and intermediate mesoderm derivatives (organs). Examples of Cluster 1-5 clonal contributions are shown (Fig. 2g).

Taken together, this analysis indicates that there were a small set of reproducible clonal fate patterns of E5.5 epiblast cells and that, within multi-lineage clones, strong biases were found among certain tissues, most notably between neural versus limb-biased multi-outcome clones (Fig. 2e-g).

### Fetal organ lineage branch point hierarchies informed by clonal relatedness of tissues

The fate of clones can differ both in terms of the type of tissues they generate and the number of cells they contribute to in each tissue (Fig. 2). We reasoned that by comparing these clone-related features between pairs of tissues, it would be possible to determine the ancestral relationships of tissues and infer the underpinning developmental processes. In this conceptual framework, tissues that have a similar clonal composition are likely to share a developmental trajectory with a ‘late’ branching point, whereas dissimilar clonal compositions may reflect an ‘early’ branching point and/or late-stage clonal expansion.

As a proof-of-principle, we first performed **c**lonal **s**imilarity **a**nalysis (CSA) using the cosine similarity between tissues as a measure of relatedness in a selection of samples that are well understood in terms of segregation during development, namely blood, brain (fore- and hindbrain, left and right), limbs (forelimbs and hindlimbs, left and right), and gonad and kidneys. We found that samples of the same tissue type were more similar in their clonal composition compared to samples originating from a different tissue (e.g., brain tissues vs limbs), likely reflecting developmental branching between the tissue types (Fig. 3a). Unfortunately, clustering was relatively noisy both on individual embryos and for all three embryos pooled probably due to insufficient numbers of informative barcodes (Fig. 3a).

**Figure 3.**
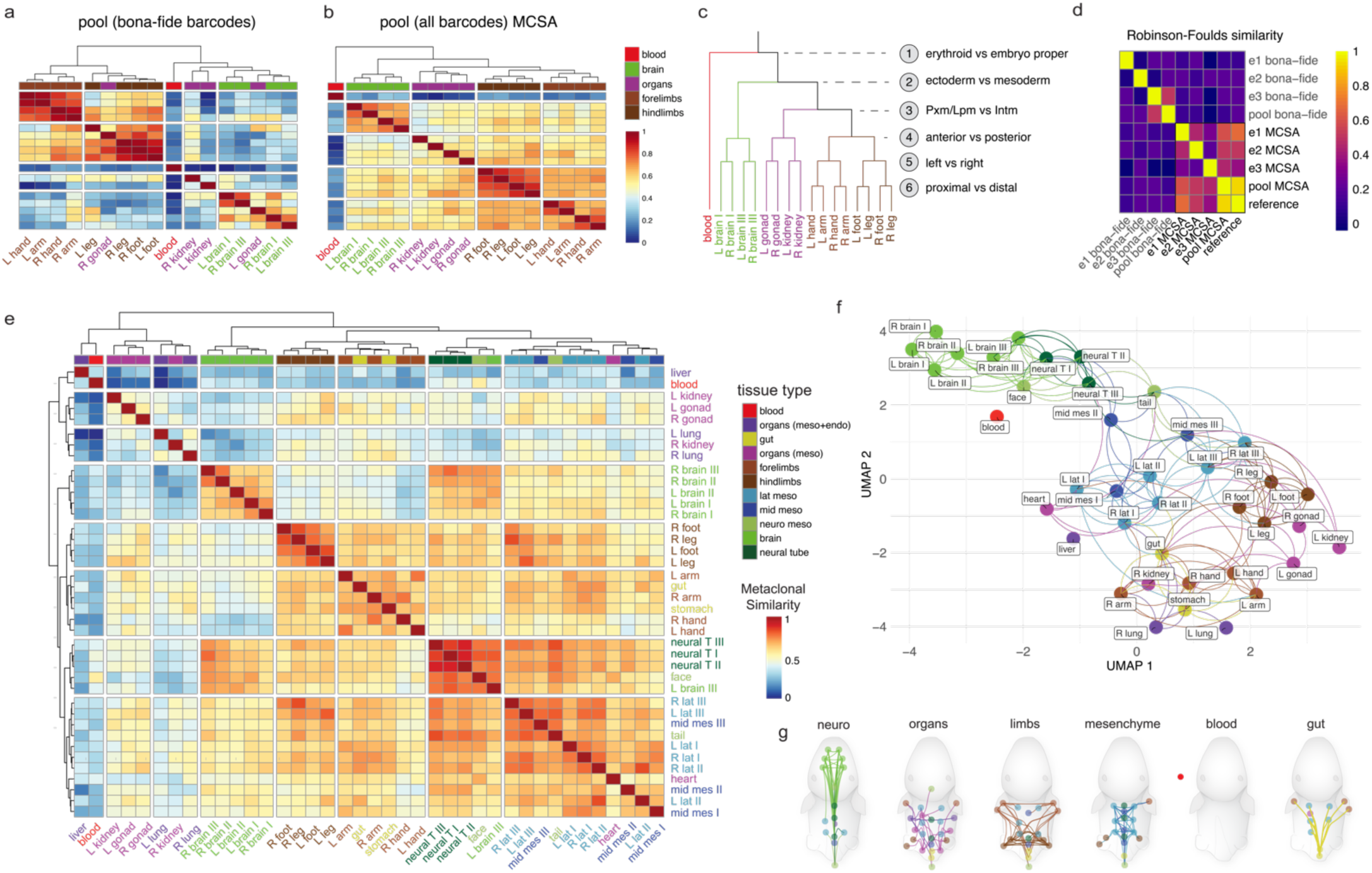
Major developmental branch points established through barcode induction at E5.5 followed by metaclonal similarity analysis at E12.5. Clustered heatmap of cosine similarity matrix computed for a selection of tissues pooled from three embryos using **a)** bona-fide and **b)** all clones (meta-clonal similarity analysis, MCSA, see text and Methods). **c)** Differentiation hierarchy of tissue development comprising 6 sequential levels of bifurcation. **d)** Robinson-Foulds similarity between hierarchies derived from bona-fide and all barcodes. **e)** MCSA of all tissues visualised as a heatmap, **f)** UMAP with lines connecting to 4 nearest neighbours with an MCS >0.75, or **g)** a pseudo-anatomical map where tissues of a given type (legend in a), were connected to their 4 nearest neighbours with an MCS >0.75. Edge colour indicates the tissue type of the outgoing node per legend in e). Data represent the aggregation of 3 biological replicates that were also assessed individually (Supplementary Fig. 4). Abbreviations: Pxm, paraxial mesoderm; Lpm, lateral plate mesoderm; Intm, intermediate mesoderm.

We noted however that the clustering of tissue types became more robust when all barcodes (including likely ‘barcode collisions’ otherwise excluded due to high probability^39^, see Methods) were incorporated for computing the cosine similarity scaled to range from 0 to 1 (Fig. 3b). Because frequent barcodes may correspond to several individual clones harbouring the same barcode by chance, i.e., what we refer to as *meta-clones*, to distinguish it from CSA we termed this approach **m**eta-**c**lonal **s**imilarity **a**nalysis (MCSA). To quantify the power of MCSA more systematically we compared branching hierarchies derived from filtered and unfiltered barcodes and from individual embryos and pooled against each other using the Robinson-Foulds similarity, a commonly used measure for the similarity between trees. In this comparison we also included a hierarchy, based on results from Fig 3b and prior knowledge from literature. This resulted in a hierarchy consisting of six sequential bifurcation points: initiated with early branching of circulating nucleated blood cells, followed by ectoderm and mesoderm specification, paraxial/lateral plate mesoderm and intermediate mesoderm, then anterior posterior, then left right, and finally proximal distal branching (Fig. 3 c-d). This analysis highlighted that hierarchies derived from unfiltered barcodes were, despite likely barcode collisions, more consistent between individual embryos and the pool and were more similar to the model hierarchy in Fig. 3c.

The fact that stringent filtering, which is the norm in *in vivo* barcoding data analyses, lead to less robust results was counterintuitive initially due to technical artifacts potentially introduced by barcode collisions. However, we speculated that a trade-off exists between the number of clones that are available to compute the similarity and the chance that some barcodes may represent more than a single clone. Next, we applied MCSA to the full data set, and utilized 3 complementary approaches to visualize the results: through a hierarchically clustered heatmap (Fig. 3e), through UMAP to project tissues (rather than individual clones as in Fig 2e) in 2D and connect MCSA-based nearest neighbours (Fig. 3f), and a pseudo-anatomical representation of tissue interconnected using edge bundling of nearest neighbours and split by a reduced set of tissue classes (Methods) (Fig. 3g). For completeness we also provide the heatmap using *bona-fide* clones in Supplementary Fig. 3a with the same ordering as in Fig 3e.

All three visualizations revealed similar groupings, consisting of neural clusters, limb clusters, mesenchymal and organ cluster, and blood clearly isolated from the other tissues. However, there were also differences. While the heatmap separated splanchnopleure/extraembryonic mesoderm from neurectoderm/mesoderm derivatives, the UMAP suggested a more global neurectoderm/mesoderm separation, with splanchnopleure tissues closely associated with paraxial mesoderm derived tissues. Due to its robustness to noise and more flexible 2D layout rather than a linear ordering, the UMAP representation is better suited for visualizing complex tissue relationships accurately.

The *in vivo* LoxCode clonal lineage tracing data highlighted a series of highly correlated groupings (Fig. 3e-f), in which tissue/organ groups have shared/similar clonal ancestry These included: (i) a brain tissues/neural tube/tail/face group that represent tissues of a neuroectoderm origin; (ii) limbs/neural tube/lateral mesoderm/medial mesoderm that represent tissues derived from mesoderm; and (iii) gonad/kidney/gut containing derivatives of intermediate/lateral mesoderm. In contrast, tissue groups that were less correlated such as liver/blood, brain/kidneys and gonad/lung were suggestive of distinct ancestry. Other examples of the least correlated samples were (i) endoderm containing tissues (liver and lungs) that were distinct from mesodermal and neuroectodermal tissues, but are associated with gut and stomach; (ii) anterior neuroectodermal derivatives (forebrain and midbrain) were clustered separately from the neural tube. The separate clustering of these two groups may be due to the distinct origin of anterior neurectoderm derived from the epiblast and the trunk neural tube derived from the neuromesodermal progenitor<u>^11^</u>; (iii) neural tissues and limbs displayed distinct clonal composition; and (iv) fore- and hindlimb displayed separate clonal origin and sidedness of clones in the limb tissues, possibly related to the separation of progenitors to contralateral lateral plate mesoderm.

### A stochastic agent-based model of embryogenesis and LoxCode barcoding

To explore potential aetiologies of the clonal similarities between samples observed in our data, we implemented a generative stochastic agent-based model of tissue development (see *Methods* and Supplementary File 1). Starting from the zygote, we simulated stochastic cell divisions^40^, LoxCode barcoding at E5.5, as well as differentiation into a selection of tissues following the lineage rules informed by known relationships between tissue types, summarized in Fig. 3c (Fig. 4a). At day E5.5, cells (i.e., agents) were assigned a LoxCode with probabilities based on a stochastic Cre recombination process^40^. Except for the first two divisions and blood progenitors (see *Methods*), division times were assumed to be 12 hours in average yielding cell numbers at E5.5 and E12.5 similar to previous estimates^3^. At day E12.5, with a simulated population size of typically over 6 million cells, clonal data from three simulated embryos (to match the actual data) was used to generate an MCSA matrix. These simulations were then run thousands of times to fit the model output to the empirical data using the evolutionary covariance adaptation strategy (CMA-ES, Fig. 4b, Supplementary Fig. 5, ^41^). A total of 33 parameters were allowed to vary including the timings *t* of fate bifurcations, *p* the probabilities to differentiate into a specific cell type at a junction, and the rate of fate determination.

**Figure 4.**
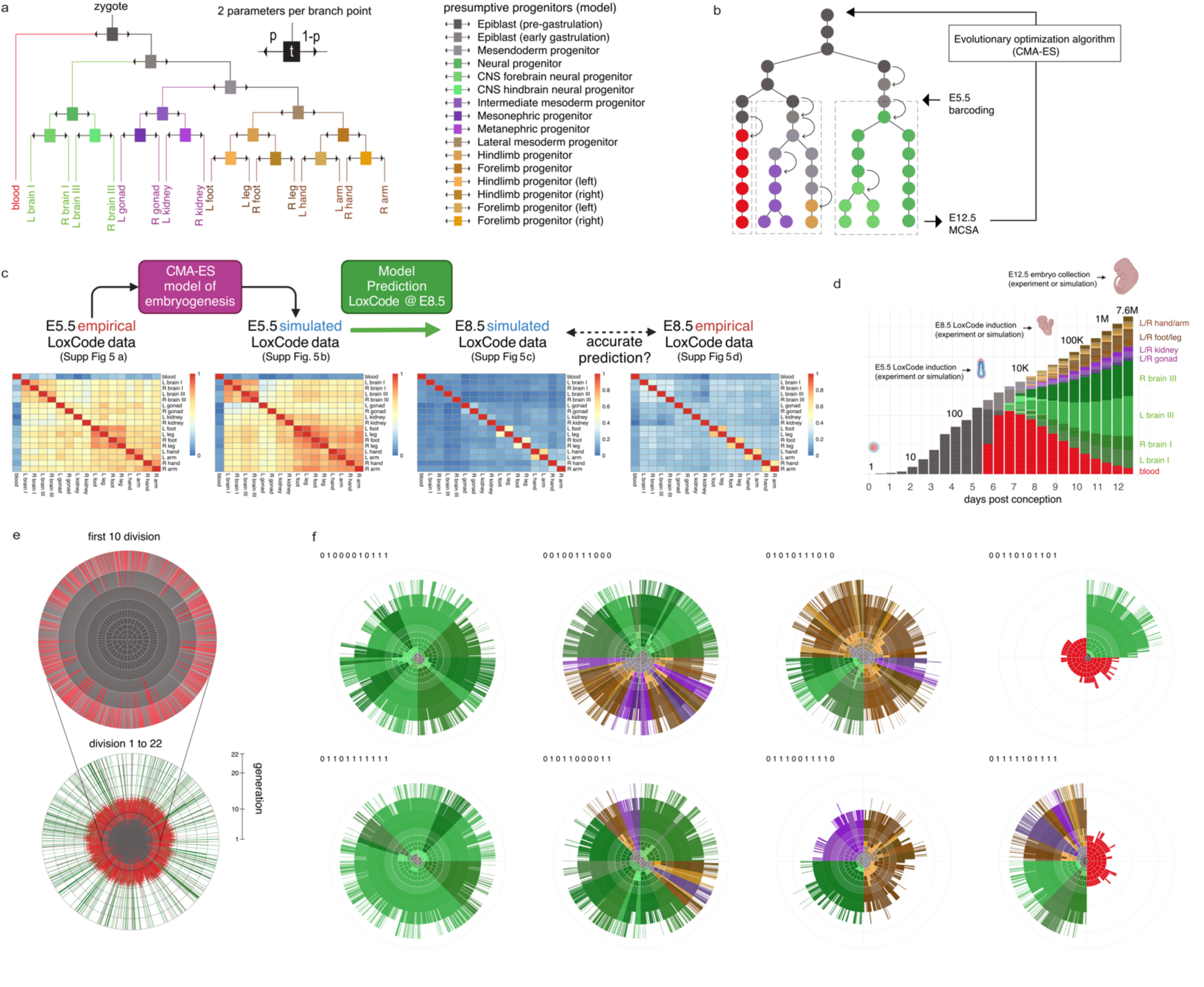
*In silico* embryogenesis. **a)** Agent-based model of tissue development incorporating differentiation rules in Fig. 3c. Model progenitor nomenclature introduced for illustrative purposes. **b)** Schematic of a realization of the model at the cellular level, including stochastic timings between divisions, differentiation and barcoding. Parameters are fitted iteratively using the CMA-ES evolutionary algorithm. Color scheme is for illustrative purposes. **c)** Comparison of empirical and model simulations for E5.5 and E8.8 barcode induction with LoxCode retrieval at E12.5. Only E5.5 data was used for fitting the model parameters. **d)** Simulated kinetics of tissue differentiation from day of conception to E12.5 with frequencies on linear and population size on logarithmic scale. Embryo icons at different stages are for illustration purposes. **e)** Nested-zoom into a simulated 22 generation pedigree, from the zygote to E12.5. Concentric circles correspond to generations (the centre is generation 0 and 1), each segment represents one cell coloured according to its cell state at division (legend, panel a). Two daughter cells are placed adjacent to the mother but on the next outer concentric circle (see *Methods*). f) Examples of simulated clonal pedigrees from single cells at generation 10, corresponding to the time of experimental LoxCode induction at ∼E5.5. The binary string next to each example indicates the ancestry of the founder cell of that clone (see Circular Pedigree Plot in Methods). Color scheme as in panel a.

Despite its simplicity and absence of spatial aspects, we found that the model could reproduce clonal similarities summarized in the MCSA heatmap (Fig. 4c), fate biases at the clonal level (Supplementary Fig. 5a, not used for model fitting), while simulating all division and differentiation events from the zygote to an approximately 7.4 million simplified E12.5 embryo (Fig. 4d). In the model, the bifurcation of the blood lineage from other embryonic lineages was inferred to occur shortly after barcoding at E5.5 followed by the ectoderm-mesoderm split approximately 24 hours later (Fig. 3d). All remaining bifurcations were completed by E9.5. Best-fit parameters from different random initialization showed a relatively narrow range indicating the parameters were well defined given the model and the data (Supplementary Fig. 5e-f).

To understand how clonal similarities between tissues would change when Cre recombination is induced at a later stage during development, we induced LoxCodes recombination at E8.5 in our simulations and analysed *in silico* embryos at E12.5. MCSA from this simulated data predicted a sharp drop in clonal similarities between tissues, except for limbs and legs, which maintained some degree of concordance (Fig. 4c). To test the validity of this prediction, we then repeated our previous experiment (Fig. 3) but administered tamoxifen at E8.5 rather than E5.5. A total of 24,568 ± 4,182 unique barcodes were detected across 3 embryos, which was consistent with the number of barcodes expected from the number of cells in E8.5 embryos. Closely aligning with model predictions (importantly the model parameters were exclusively fitted to the E5.5 MCSA data) and consistent with known biology, the MSCA heatmap revealed an equally striking loss of clonal relationships compared to E5.5, with the exceptions of limbs and legs (Fig. 4c). Interestingly, and matching the E5.5 analysis, general resemblance between empirical and simulations were maintained beyond MCSA at a finer-grained clonal level, when filtering on large clones (frequency > 0.1%) detected in more than one tissues (Supplementary Fig. 5c). One aspect where the model and empirical data differed was the increased barcode sharing between tissues in the real world data. A possible explanation for this observation could be circulating blood cells, an aspect currently not incorporated into our model. Besides serving as an important test case for our generative agent-based model of embryogenesis, barcoding at a later time point with an approximately 50-100-fold increase in number of cells per embryo relative to E5.5 demonstrated the scalability of the LoxCode technology towards larger population sizes.

Finally, to visualize full pedigrees generated in our simulations, which due to the large number of cells was impractical with ordinary hierarchical trees, we positioned cells on concentric circles according to generation, ordered within and between circles such that mother cells neighbour their daughter cells, and coloured according to the cell’s state at division (Fig. 4e). Using a similar approach, we then visualized E5.5 clones (Fig. 4f), illustrating how fate diversity is generated at the clonal level. In summary, fate biases, in this model, were caused by early differentiation events that occur shortly after barcoding, while similarities in clonal frequencies between tissues arose as a consequence of shared differentiation paths.

Overall, the combination of stochastic agent-based modelling with MCSA suggested that a putative hierarchical model of developmental branch points and timings can be inferred from a single snapshot of barcode induction. In the future, incorporation of clonal data at additional time points could be used to refine the model and empower the discovery of so far uncharacterized branch points for elucidating the precise order and timing of each bifurcation.

### Determination of cell type bias and clonal relationships at a single cell level

Bulk barcode analysis has the advantage of examining millions of cells, unlike single-cell analysis, which typically studies thousands due to cost and throughput limitations. Bulk analysis therefore allows for a more comprehensive view of clones, including the detection of rare clones that might be missed with single-cell omics analysis. However, bulk barcode analysis does not reveal which cell types contain any given barcode within a tissue. As the LoxCode construct is by design actively transcribed under the endogenous Rosa26 promoter, combined with single-cell RNA-sequencing, it enables simultaneous determination of cell type and clonal origin. We dissected an E12.5 embryo barcoded at E5.5 into organs/tissues as outlined in Fig 1b. From this, single cell suspensions were generated, multiplexed using MultiSeq^42^ (Methods, Supplementary Fig. 7a), and processed on a droplet-based scRNAseq platform to capture ∼30,000 cells. Following quality control (Methods), 33 cell types (Fig. 5a) across all tissues (Fig. 5b, Supplementary Fig. 7b) were annotated (Methods, Supplementary File 1).

**Figure 5.**
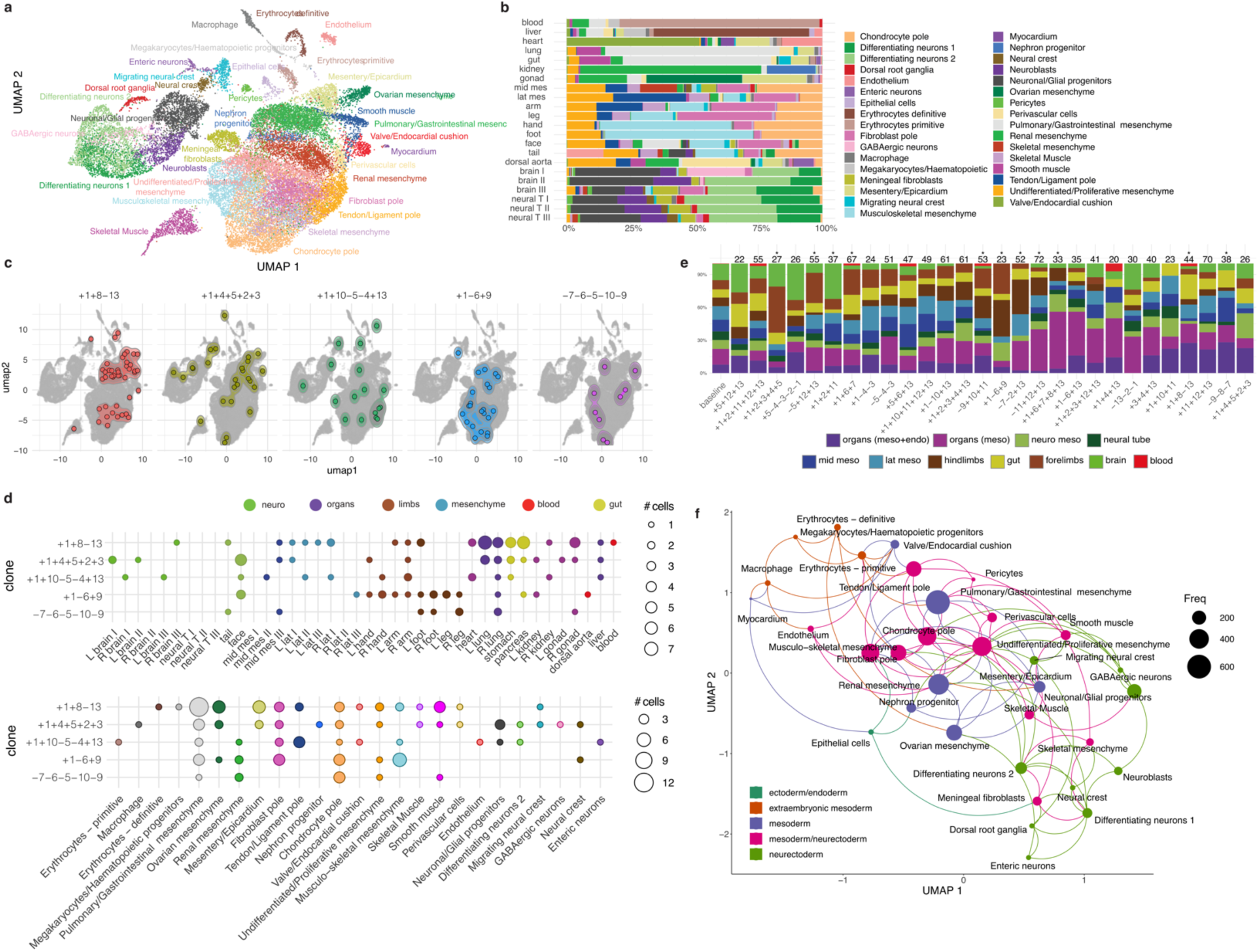
Single cell readout of epiblast fate bias and clonal relationships. **a)** UMAP of scRNA-seq derived from ∼30,000 cells and annotated cell types, with **b)** their distribution across tissues (left and right and mesenchymal tissues pooled). **c)** Examples of single cell bona-fide clonal output corresponding to the indicated LoxCodes. Distribution of single cells per detected LoxCode in c) according to **d)** tissue (top) or cell type (bottom). **e)** Tissue type distribution of meta-clones with at least 20 cells and ordered using 1D UMAP. * indicates barcodes where distribution significantly deviates from a base-line distribution (χ2 test, adj. p-value < 0.05 using Benjamini-Hochberg correction). **f)** scMCSA UMAP representation of cell types from b), with lines connecting each cell type’s 3 nearest neighbours. Dot size is proportional to the cell number in each cluster, and colouring is according to broad cell type categories as indicated in the legend.

Barcodes were then selectively amplified from the single-cell full length cDNA libraries prior to fragmentation of the cDNA library, and both were sequenced separately on a NextSeq (for cDNA) and MiSeq (for LoxCodes) and mapped *in silico* by virtue of their shared 10X cell barcode. Overall LoxCode capture efficiency was 14.6% (4659 cells), but we noted recovery as low as 6% in neuronal tissues. After filtering out frequent LoxCodes (Methods), we recovered ∼505 cells (10.8% amongst cells with a LoxCode detected), spanning 33 bona-fide clones with at least two cells, of which 15 contained >8 cells (Supplementary Fig.7).

From these 15 clones we selected 5 examples with distinct fate patterns (Fig. 5c). Cellular distribution of these clones indicated contribution to a large range of cell types, but also biased distributions across tissues (p-value < 0.0001) and cell types (p-value < 0.0001) (Fig. 5d). At the tissue level, for instance clone +1+8-13 was enriched in lungs, gut and mesenchymal tissues, clone +1-6+9 was prominent in fore- and hindlimbs, with low detection frequency across all organs. Clone +1-6+9, +1+4+5+2+3, and +1+10-5-4-13 were detected in face, but only for the latter two were present in brain. At the transcriptional level, all 5 clones had representatives in the pulmonary/gastrointestinal mesenchyme and chondrocytes. Clone +1+4+5+2+3 contributed to a diverse set of neuronal cells, while clone +1-6+9 differentiated mostly to musculoskeletal mesenchyme, chondrocyte and fibroblast cells consistent with its limb-biased nature.

Overall, despite the relatively low number of clones that could be assessed, heterogeneity in transcriptional fates was clearly evident. To explore if meta-clones could provide additional insights, we repeated the analysis for barcodes for which we were less confident that they originated from a single cell. First, we confirmed that tissue type contribution deviated from the population baseline (Fig. 5e). Bias towards forelimbs (light-brown color) or hindlimbs (dark-brown color) was apparent for meta-clone +1+2+3+4+5. Relative to baseline, meta-clone +1+6+7+8+13 showed increased presence in mesodermal organs, coinciding with a low detection in endoderm organs such as lungs, liver and gut. Meta-clone +1+4+13 was not detected in limbs but in blood (1 cell), although this was not statistically significant. Interestingly most meta-clones were detected in gut. Among meta-clones we identified 4 additional candidates with apparent transcriptional bias (Supplementary Fig. 8a-c). While we cannot directly comment on the fate of these clones (i.e., they could represent the combined fate of two or more epiblast cells), their heterogeneity adds support to the previous observations (Fig 3) that most E5.5 epiblast clones are biased in their fate.

Global relationships between cell types were analysed using a single cell version of MCSA (scMCSA). In this approach meta-clonal similarities between transcriptional cluster (instead of tissues as in Fig. 3f) were computed and projected onto 2D using UMAP (Fig. 5f). Cell types broadly clustered into 1) haematopoietic and cardiac, 2) neuronal, and 3) mesenchymal types. Strong relatedness between haematopoietic and cardiac tissues was consistent with the notion that mesodermal cells that contribute to both tissues, are the first cells to be allocated during gastrulation, and might share common epiblast ancestors. These results complement the findings in Fig. 3f which showed a clear distance between blood, neuronal and mesenchymal tissues at the population level.

### Asymmetric contribution of epiblast clones to gonads and kidneys at the single cell level

Measuring similarities in clonal compositions between different tissues (Fig. 3-5) revealed several clusters of strong and expected clonal relationships, notably brain and neural tissue/cell type inter-relatedness, and limb tissues. Surprisingly, however clonal similarities between kidney and gonad were found to be relatively weak, which was unexpected considering kidney and gonads emerge from the urogenital ridge at E9.5, a time much later developmentally compared to our LoxCode induction time of E5.5. To investigate how this result could be explained, we repeated the experiment of Fig. 5 with a focus on cells exclusively from both left and right kidney and gonads of an E12.5 embryo after barcode induction at E5.5.

A diverse array of cell types was present across left and right kidney and gonad (Fig 6a), with LoxCode detection efficiency in gonad and kidney of 30% and 38% respectively. A total of 108 unique complex barcodes were detected (corresponding to 1014 cells or 6.4% of cells in which a LoxCode was detected), of which 51 clones had more than one cell, and 18 clones with >8 cells. Clones were then positioned according to their left/right or kidney vs gonad organ bias (Fig 6e). Strikingly, most clones, especially large clones, were significantly biased in their contribution to an organ or L/R symmetry, or both, with a trend towards left kidney bias. The distribution of cells from four example clones is shown in Fig 6f where kidney vs gonad and/or left vs right asymmetry was apparent. Therefore, despite kidney and gonad sharing a common developmental trajectory until very late in embryogenesis, epiblast contribution to these organs was largely asymmetric.

**Figure 6.**
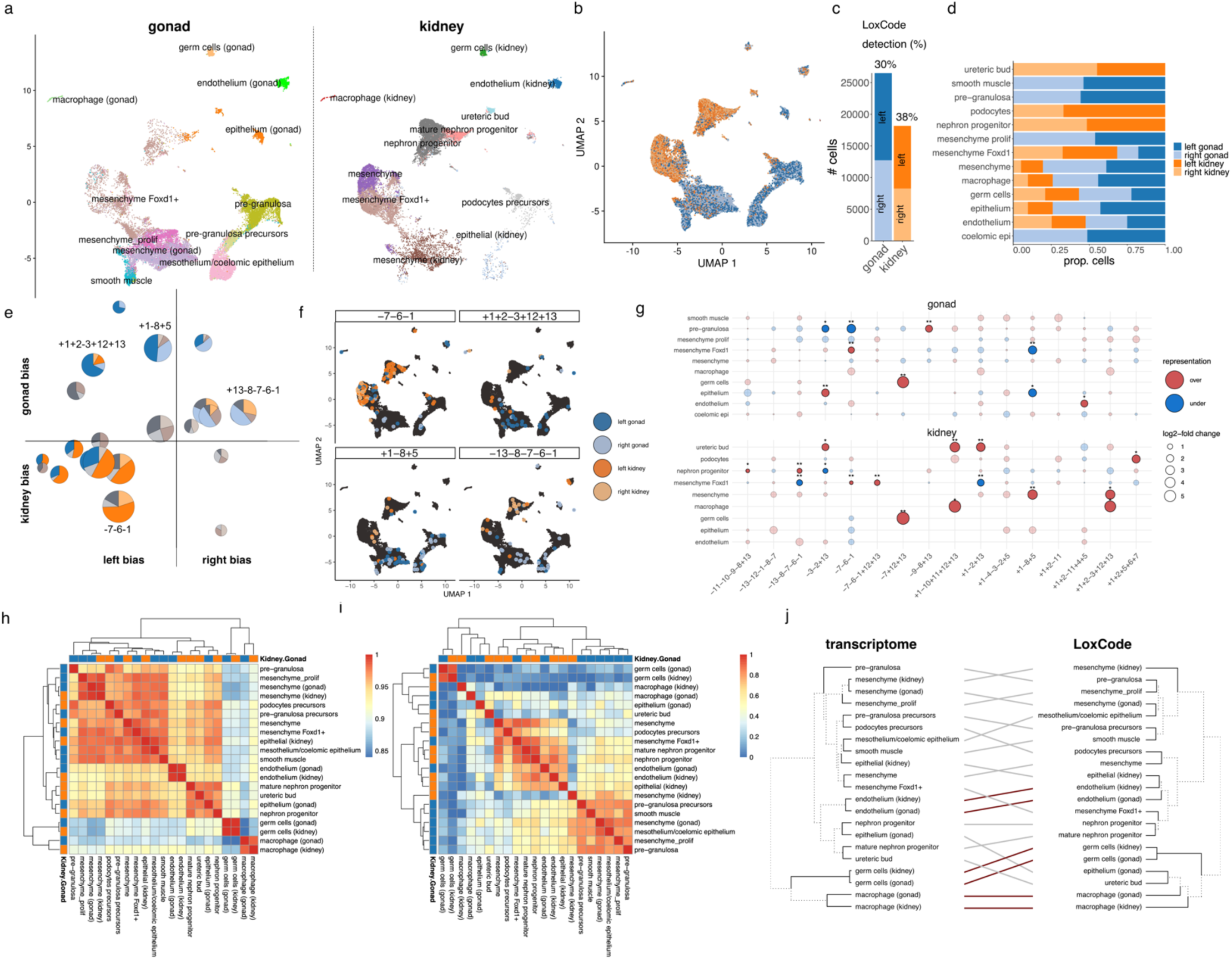
Asymmetric contribution of epiblast clones to gonads and kidneys. **a**) Single-cell RNA sequencing data of >40K cells from E12.5 embryonic gonad (left: 13769, right: 12735) and kidney (left: 9896, right: 8264) induced at E5.5. UMAP projection was performed on the combined data set, then split by tissue. **b**) UMAP projection coloured according to tissue of origin. **c**) Number of cells from left and right gonad and kidney and LoxCode detection frequencies. **d**) Proportions of left and right gonad and kidney cells across transcriptional clusters. **e)** Gonad-kidney and left-right bias per epiblast clone. Colour indicates significant bias (G-test, adj. p-value<0.05). Size is proportional to log2 of the number of cells per clones. **f**) Individual examples clones annotated in panel e) overlaid onto scRNA UMAP. **g**) Gonad-kidney cell-type bias (* p-value<0.05, ** adj p-value <0.05). Colour indicates whether the cell type is over- or underrepresented, the log2-fold change is proportional to the dot size. **h**) Transcriptional similarity between transcriptional clusters in gonad and kidney. **i**) Clonal similarity (scCSA) between transcriptional clusters in gonad and kidney. Color code as in panel d). j) Dendrograms for both transcriptional and ontogenic similarity are shown, with cell types linked and coloured according to inter-dendrogram concordance. Links are coloured according to relatively high transcriptional-clonal concordance (red, inter-leaf distance in both dendrograms ≤2) or low (grey, distance >2 in at least one dendrogram).

Next, we analysed clonal origin of cell types across both organs. We enumerated how many cells of each type were produced by each clone with at least 10 cells (Supp Fig 9a). Then we compared this to the overall distribution of cell types in the entire dataset. This allowed us to quantify if a clone generated more or fewer cells of a particular type than what would be expected based on the overall distribution (Fig 6g). Of the 16 clones, 8 generated cell types that were statistically over- or under-represented. There were some notable examples. Clone -7+12+13 was strongly enriched in primordial germ cells (PGCs) and interestingly was detected at similar frequencies not only in left and right gonad but also kidneys. This is in line with an early branching of the germ lineage from mesenchymal and endothelial tissues and migration from mesentery to genital ridges at around E10.5. Clone +1-8+5 preferentially contributed to kidney mesenchyme but not gonadal Foxd1 mesenchyme. Lastly kidney biased clone +1-10+11+12+13 preferentially gave rise to macrophage and ureteric bud. These relationships do not immediately fit with prior knowledge, and future studies are needed to determine the robustness of these fate patterns.

A long-standing question in developmental biology is whether cell types or tissues with similar gene expression profiles (i.e., transcriptionally related) are likely to be ontogenically related (i.e. clonally related). To test this in our system, we separately assessed the transcriptomic similarity between cell types (Fig 6h) and clonal similarity using scCSA (Fig 6i, Supp Fig 8b). Transcriptionally defined cell types clustered broadly into germ cell, macrophage, endothelial, nephron, and mesenchymal/epithelial clusters, and this was largely independent of organ. Ontogenically, however, cell types largely clustered according to their organ of residence with some exceptions (for germ cells and macrophages, which aligns with their known migratory behaviour). To assess more systematically how ‘concordant’ the transcriptomic and ontogenic relationships were between the same cell types, we untangled and compared the respective hierarchical clustering dendrograms and coloured connections according to inter-leaf distance (Fig 6j, red: ≤2 inter-leaf distance in both dendrograms, grey: >2 inter-leaf distance in at least one dendrogram). This result highlights that transcriptomic-ontogenic concordance is cell type dependent and the exception rather than the rule for the majority of E5.5 epiblast clones contributing to kidney and gonadal tissues.

The similarities in clonal compositions of germ cells and macrophages between left and right sides, as well as between kidney and gonad aligns with our understanding of the migratory nature of these cell types and their early specification in development. The observation that other clones show significant imbalances, with biases towards either left vs. right or kidney vs. gonad, was unexpected. The left-right imbalances in clonal contribution challenge the notion of strictly independent and symmetric development of these bilateral structures. This hints at potential differences in the proliferation rates or migratory behaviours of progenitor cells on either side of the embryo. The kidney-gonad bias in clonal contribution is particularly intriguing, as it suggests that some level of fate restriction between kidney and gonad lineages may occur earlier than the segregation of cell types in the urogenital ridge at E9.5, although local expansion cannot be ruled out. These findings raise important questions about the timing and mechanisms of cell fate decisions in early urogenital development. They suggest that the process may be more complex and occur earlier than previously understood, with possible implications for our understanding of congenital anomalies affecting these organ systems. Further investigation into the molecular mechanisms driving these early clonal biases could provide valuable insights into the fundamental processes of organ specification and the establishment of left-right asymmetry in the urogenital system.

In sum, through LoxCode barcoding of the mouse epiblast we have uncovered the patterns of the clonal contribution to tissues and cell types of the late organogenesis-stage embryo and the inter-relationship of fetal tissues and organs.

## Discussion

Using the LoxCode mouse model presented herein, we find pre-gastrulation (E5.5) epiblast cells are biased in their clonal contributions across tissues and cell types in the germ layer derivatives: notably towards either blood, ectoderm lineages, mesenchymal tissues, or limbs, likely reflecting developmental branch points during development. How do these developmental branch points occur? Our simulated clonal pedigrees infer that fate restriction commences around E5.5 and completes by E9.5. However, presently, our results do not allow discrimination between several non-mutually exclusive aetiologies: epiblast molecular fate imprinting^43^, multi-lineage epiblasts that, by virtue of their physical location, receive specific signals to later restrict their fate^44^, lineage specific timing of allocation to specific cell/tissue type during development^45^, and late-stage clonal expansion of the specified progenitor or the descendant tissue/cell types^46^.

The notion of fate bias is well-established within the field of embryology, particularly during the initial phases of development stages^47,48^. In contrast to a binary approach that classifies cells according to their capacity to differentiate into a specific assortment of mature types (e.g., bi-potent, tri-potent), a framework of fate bias presents a more refined evaluation of relative contributions. This concept is key to the latest generation of revised models of hematopoiesis^49^ that, instead of rigid binary bifurcations, favour a ’continuous’ model and acknowledge the existence of multi- and oligopotent progenitors exhibiting varying degrees of lineage-bias early in the stem and progenitor cell hierarchy. Throughout development, each stage is characterized not only by the current cellular state but also by its origins and future differentiation potential. The LoxCode mouse model and the accompanying analytical tools introduced here now offer a higher-throughput approach to quantify these fate biases and their progression not only for embryogenesis, but also in other systems like hematopoiesis, immunology and cancer.

The key features of the LoxCode mouse that empower this approach include barcode generation that is expeditious (within 3-12 hrs of Cre-ER^T2^ activity, the majority of barcodes recombine at least once), using low-dose tamoxifen induction, and efficient (a stable identity and high diversity of >30 billion LoxCodes, >2000 size-stable barcodes resistant to further Cre recombination), with barcodes detectable through PCR or single cell RNA-seq. This model allows a significantly higher throughput compared to existing Cre LoxP based barcoding^23,24^ by using short read sequencing for barcode read-out.

An important next challenge will be to connect fate bias across different stages of development. Pioneering technologies including evolvable hgRNA barcodes^19^ already address this question but are currently hampered by i) the non-randomness of NHEJ outcomes^50^ which limits the diversity of founder cells, and ii) the lack of rapidly inducible Cas9 mouse models. Crossing the LoxCode mice with the MARC1 system or CARLIN^20^ or other CRISPR based barcoding technologies could represent an efficacious paragidm to precisely map fate bias kinetics over time, by combining the best of Cre LoxP (i.e., efficiency, temporal control, high baseline diversity) with Cas9 based approaches (i.e., accumulation of mutations, exponential increase in diversity as a clone expands). In addition, constructs using alternative site-specific recombinases or integrase (e.g., Flp, Bxb1), or emerging technologies like transcriptional cell history recorders^26^ could provide additional levels of resolution and the recording of molecular correlates of branching.

While we are far away from replicating the notable work in *C.elegans*^51^ for mouse development^52^, the current study provides a comprehensive view into clonal fate heterogeneity of E5.5 epiblast in murine embryos and demonstrates the LoxCode technology as a powerful tool to characterize clonal fate at exceptional depth and higher throughput.

## Methods

### LoxCode cassette restriction cloning

Following multiple unsuccessful attempts to synthesize the LoxCode cassette through commercial service providers, presumably due to the palindromic characteristics of LoxP repeats, we pursued an alternative approach involving the assembly of the LoxCode cassette via sequential restriction cloning. To this end, we designed and procured two partially reverse complementary 5’ phosphorylated 90 base pair oligonucleotide libraries (Integrated DNA Technologies, duplex oligos), which encompassed two LoxP sites in opposing orientations (underscored), an intervening 8-base sequence consisting of mixed bases (Ns), and flanking regions comprising 4 mixed bases. Additionally, each oligonucleotide contained a 5-base extension at the 5’ terminus, facilitating the formation of sticky 5’-AATT-3’ overhangs upon annealing, thereby rendering the duplex oligo compatible with ligation into EcoRI-digested plasmids. The nucleotide base adjacent to the sticky end was selectively designated as cytosine (C) and adenine (A) (highlighted in bold), ensuring that the EcoRI (GAATTC) recognition site would be reconstituted solely at one extremity following ligation, thereby permitting sequential integrations, i.e.

5’-AATT**C**NNNNATAACTTCGTATAATGTATGCTATACGAAGTTATNNNNNNNNATAACTTCGTATAGCATACATTATACGAAGTTATNNNN**A**-3’ 3’-**G**NNNNTATTGAAGCATATTACATACGATATGCTTCAATANNNNNNNNTATTGAAGCATATCGTATGTAATATGCTTCAATANNNN**T**TTAA-5’

Subsequently, the pUC18 plasmid was subjected to digestion with the EcoRI-HF restriction enzyme (New England Biolabs), followed by a standard ligation reaction. In order to minimize the likelihood of background recombination in bacterial cells, Sure 2 (Agilent) competent cells were transformed with the ligation product and cultured at reduced temperatures (27°C). Individual colonies were then screened for integration events and orientation via restriction digest analysis. Plasmids isolated from successful colonies, exhibiting the reconstituted EcoRI site at the 3’ terminus of the cassette, underwent a subsequent round of EcoRI digestion for further cloning. This procedure was iteratively performed seven times, ultimately yielding a nucleotide sequence encompassing a total of 14 LoxP sites and 13 random barcode elements, with either 8 (odd position, NNNNNNNN) or 14 base pairs (even position, NNNNGAATTTNNNN).

ATAACTTCGTATAATGTATGCTATACGAAGTTAT**ACTCCGCA**<u>ATAACTTCGTATAGCATACATTATACGAAGTTAT</u>**TCCAGAATTTGTAT**ATAACTTCGTATAATGTATGCT ATACGAAGTTAT**ACATCCAC**<u>ATAACTTCGTATAGCATACATTATACGAAGTTAT</u>**AAAGGAATTTCTCC**ATAACTTCGTATAATGTATGCTATACGAAGTTAT**ATTTCCTC**<u>AT</u> <u>AACTTCGTATAGCATACATTATACGAAGTTAT</u>**GCCCGAATTTTTTC**ATAACTTCGTATAATGTATGCTATACGAAGTTAT**GCTACTGG**<u>ATAACTTCGTATAGCATACATTAT</u> <u>ACGAAGTTAT</u>**ATGAGAATTTATGG**ATAACTTCGTATAATGTATGCTATACGAAGTTAT**AACTAGAA**<u>ATAACTTCGTATAGCATACATTATACGAAGTTAT</u>**TGCAGAATTTCC TC**ATAACTTCGTATAATGTATGCTATACGAAGTTAT**CGACACTT**<u>ATAACTTCGTATAGCATACATTATACGAAGTTAT</u>**AACGGAATTTTCAA**ATAACTTCGTATAATGTATG CTATACGAAGTTAT**CGTGTTTG**<u>ATAACTTCGTATAGCATACATTATACGAAGTTAT</u>

### LoxCode mouse generation

The LoxCode mouse line was generated following a previously established protocol^53^. In brief, double strand breaks were introduced at the *Rosa26* locus in C57BL/6J oocytes via microinjection of Cas9-gRNA ribonucleoproteins (gRNA: 5’-CTCCAGTCTTTCTAGAAGAT -3’), co-injected with a circular HDR donor vector (pBlueScriptIISK backbone) containing a splice acceptor, the LoxCode sequence, and a polyA signal flanked by 1000-1500bp Rosa26 specific homology arms. Oocytes were transplanted into pseudopregnant females, and 2 out of 26 viable pups displayed the expected integration via PCR and gel electrophoresis. Sanger sequencing was performed to confirm the integrity of the LoxCode sequence. The LoxCode line was then bred to homozygosity on a C57BL/6 background.

### LoxCode theoretical and practical diversity

The LoxCode construct consists of 13 barcode elements, with 7 at odd positions (e.g., 1, 3, 5, 7, etc.) and 6 at even positions (e.g., 2, 4, …, 12). Recombination results in inversions and excisions that can alter the locations and orientations of individual barcode elements. However, by design, odd and even barcode elements always maintain their respective positions. We have previously demonstrated, through mathematical reasoning, that all barcodes can be generated via a series of inversions and excisions^29^. As a result, the theoretical diversity of a cassette containing n odd barcode elements and m = n - 1 even barcode elements can be calculated by enumerating all permutations and considering both forward and reverse orientations:

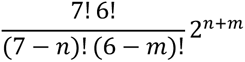

Summing this formula for barcodes of size 1, 3, 5, 7, 9, and 13 yields a total of 30,204,722,030, or over 30 billion theoretical barcodes. It is important to note that barcodes with 11 elements are not expected to be generated due to the DNA size constraint of Cre recombination, although we detected them at low frequency.

The barcode diversity generated in an experiment depends on the number of recombination events (see Table 1), which are influenced by Cre activity. Insufficient Cre results in limited barcode diversity, while excessive Cre activity leads to an accumulation of less diverse barcodes with fewer elements. Theoretically, all barcodes can be reached with 15 recombination events (see Table 1); however, barcodes with fewer elements typically achieve full diversity sooner. For example, generating all possible 201,600 five-element barcodes requires nine recombinations, while five recombinations are sufficient to produce all possible 3-element barcodes.

The necessary barcode diversity for accurate lineage tracing is determined by the number of clones tracked simultaneously. Generally, we recommend maintaining a 100-fold excess of barcodes relative to founding cells, which for the E5.5 is approximately 20,000 barcodes or at least three recombination events per cell. We typically achieve a mean of around four recombinations with a maximum of eight or nine. Across all our experiments to date (including those outside this work), we have identified over 125,000 unique barcodes, with a maximum of 11 recombination events (e.g., -3+12-9-10+7+4-11+8+5).

In addition to theoretical barcode diversity, it is crucial to consider uneven probabilities. Some barcodes are much more likely to be generated than others, which reduces effective diversity as well as the proportion of cells that can be confidently tracked within a given population.

### 4-OH-Tamoxifen preparation and administration

4-Hydroxytamoxifen Z-isomer (4-OHT, Sigma) was dissolved in 100% EtOH, diluted 1:1 with Kolliphor and stored in aliquots at –20 °C. On the day of the injection the dissolved 4-OHT was further diluted in PBS (50 µg/100 µl) and intravenously injected according to a procedure developed in Chevalier *et al.^54^*. Pregnant females received 50-100 µg of 4-OHT via tail vein injection 5 days after a plug was detected. Injected mice were separated from untreated females and monitored regularly. All animal experiments were approved by The Walter and Eliza Hall Institute animal ethics committee.

### Embryo dissections

*Rosa26CreERT2/Rosa26CreERT2* and *LoxCode/LoxCode* or *LoxCode/+* mice were set up and plug checked daily. Noon of the day of plug was referenced as 0.5 embryonic day (E0.5). Embryos were dissected in their yolk sac at E12.5 in PBS with 7% Foetal Calf Serum. The yolk sac was used as a genotyping sample when necessary (*LoxCode/+* mother). Embryos were staged using morphological criteria and assessed for normal development. The head was set aside and spilling blood was collected (blood sample). Heart, right and left lung lobes, liver, stomach (with spleen and pancreatic buds attached), intestine, left and right kidneys, left and right gonads, the region containing the dorsal aorta at the level of the aorta-gonad-mesonephros (AGM) area were dissected next. The rest of the body was cut transversely below the forelimb buds. From the anterior segment (forelimb level or level 1), left and right forelimb buds (termed L hand, R hand) were removed, followed by the left and right arms (L arm, R arm). The neural tube was removed (neural T I segment). Mesenchymal tissues were dissected as medial (left and right medial mesenchyme: L mid mes I and R mid mes I, encompassing somites) and lateral (left and right lateral mesenchyme: L lat I and R lat I, encompassing everything lateral/medial beyond the somites). The rest of the body was then cut just above the hindlimb buds level (between limb buds level or level 2), and corresponding neurectodermal or mesenchymal segments (neural T II, L mid mes II, R mid mes II, L lat II, R lat II) were dissected as above. The tail was removed and the third axial level (hindbud level or level 3) was dissected as per - level 1 (left and right hindlimbs (termed L foot, R foot), left and right hips (termed L leg, R leg), neural T III, L mid mes III, R mid mes III, L lat III, R lat III). The facial mesenchyme (face) was removed from the head, leaving the neurectoderm behind. Forebrain (brain I), midbrain (brain II) and hindbrain (brain III) segments were dissected according to morphological landmarks and halved along the midline (L brain I, R brain I, L brain II, R brain II, L brain III, R brain III). This resulted in a total of 43 embryonic samples. At the E12.5 stage, the primordia of the spleen and pancreas develop on the outer surface of the stomach, making it challenging to dissect them without cross-contamination. Consequently, these tissues were either co-dissected alongside the stomach or pooled together for subsequent analysis.

### Tissue lysis, PCR LoxCode amplification and sequencing library preparation

Tissue samples were lysed in DirectPCR lysis reagent (cell) (Viagen Biotech), and the LoxCode sequence amplified using a 3-round PCR. The first round PCR (20 cycles) uses LoxCode specific primers that efficiently amplify the LoxCode sequence from genomic DNA. As an internal quality control, for each sample, two PCR reactions were performed.

FWD primer: TCTAGAGGATCCCCGGGTACCGA

REV Primers: TGATCCGCGCCTGGATGAAT

In the second round (5 cycles), overhang primers add a partial Illumina Read 1 and Read 2 sequencing primer site, as well as a stagger between the sequencing primers and the LoxCode sequence in order to increase diversity.

FWD Primers

TCGTCGGCAGCGTCAGATGTGTATAAGAGACAG TCTAGAGGATCCCCGGGTACCGA TCGTCGGCAGCGTCAGATGTGTATAAGAGACAG N TCTAGAGGATCCCCGGGTACCGA TCGTCGGCAGCGTCAGATGTGTATAAGAGACAG NN TCTAGAGGATCCCCGGGTACCGA TCGTCGGCAGCGTCAGATGTGTATAAGAGACAG NNN TCTAGAGGATCCCCGGGTACCGA

REV Primers

GTCTCGTGGGCTCGGAGATGTGTATAAGAGACAG GCCTGGATGAATTCGTGTATAACTTCG GTCTCGTGGGCTCGGAGATGTGTATAAGAGACAG N GCCTGGATGAATTCGTGTATAACTTCG GTCTCGTGGGCTCGGAGATGTGTATAAGAGACAG NN GCCTGGATGAATTCGTGTATAACTTCG GTCTCGTGGGCTCGGAGATGTGTATAAGAGACAG NNN GCCTGGATGAATTCGTGTATAACTTCG

The third round (5 cycles) completes the sequencing primers and adds the P5 and P7 sequencing adaptors as well as introduces a unique sample index.

FWD primer:

AATGATACGGCGACCACCGAGATCTACAC[i5]TCGTCGGCAGCGTC

REV Primers

CAAGCAGAAGACGGCATACGAGAT[i7]GTCTCGTGGGCTCGG

To remove unused primers, primer dimers, and to counteract preferential amplification of shorter LoxCodes, amplicons were purified using SPRI beads at a 0.9:1 ratio between each round. All PCR reactions are carried out in a 20 µl volume, using Kapa Hifi Hot-Start Mastermix (Merck), using the following program 95 °C 2’, N times (98 °C 20’’, 65 °C 20’’, 72 °C 30’’), 72 °C 5’, 4 °C hold, where N varied depending on the PCR round. Sequencing libraries were pooled at equal volumes and sequenced on the Illumina MiSeq 600 cycle kit, using for Read 1 and Read 2 354 and 264 cycles respectively.

### Dissociation of tissues

For 10X analysis, dissected tissues (which includes all tissues except the blood sample) were dissociated for 30 mins at 37 °C in 200ul of Liberase^TM^ (Roche, 100ug/ml in PBS). 2 mL of cold PBS (without ca/Mg) + 7% FCS + 2.5mM EDTA (FE buffer) was added to the tubes and carefully removed to rinse the enzyme off. 1 mL of fresh FE buffer was added. After 15-20 mins on ice, each tissue was mechanically dissociated and filtered on a 40 μm sieve. Cells were pelleted (5 mins, 1500 rpm, 4 °C), counted with an automated cell counter (Invitrogen). 100k cells from each tissue was processed for MultiSeq labelling following the protocol described in McGinnis *et* al.^42^. For each barcode oligo (5’-CCTTGGCACCCGAGAATTCCA**NNNNNNNN**A_30_-3’), the oligo and anchor LMO (5’-TGGAATTCTCGGGTGCCAAGGgtaacgatccagctgtcact-Lipid-3’) were diluted to 2 µM in 1:1 molar ratio in PBS. The co-anchor LMO (5’-Lipid-AGTGACAGCTGGATCGTTAC-3‘) was diluted to 2 µM in PBS. Cells were resuspended in 200 µL PBS, 23 µl of anchor-barcode mixture added, and incubated on ice for 5 minutes. Then 23 µl of co-anchor solution was added and kept on ice. After an additional 5 minutes, labelling was quenched by addition of 1 mL of 1% BSA in PBS (ice cold). In all subsequent steps, cells were kept in 1% BSA on ice. After MultiSeq labelling, cells were resuspended in FACS buffer (PBS (without Ca/Mg) + 7% FCS) and labelled with 7AAD. Live cells were sorted on an Aria II cell sorter. 10K cells of each tissue were used to create 4 pools for 4 10X lanes: Pool 1 (blood, heart, right and left lung, liver, stomach, intestine, right and left kidney, right and left gonad, face), Pool 2 (dorsal aorta, L hand, R hand, L arm, R arm, L foot, R foot, L leg, R leg, L lat I, L lat II, face), Pool 3 (L lat III, R lat I, R lat II, R lat III, L mid mes I, L mid mes II, L mid mes III, R mid mes I, R mid mes II, R mid mes III, L brain I, face), Pool 4 (R brain I, L brain II, R brain II, L brain III, R brain III, neural T I, neural T II, neural T III, tail, face). The “Face” sample was seeded into each pool to allow for batch effect correction. Each pool was centrifuged (5 mins, 1500 rpm, 4C), resuspended in 32ul of PBS + 1% Bovine Serum Albumin and counted before loading on Chromium controller (10X Genomics) for scRNAseq. Sequencing libraries were prepared according to manufacturer’s recommendations and sequenced on the Illumina platform (NextSeq 75 cycle kit).

### Target amplification (scRNAseq)

The LoxCode sequence was amplified from full length cDNA using a combination of PCR, biotin capture, restriction digest and adaptor ligation. In a first round, transcripts contained the LoxCode sequence were enriched from full-length cDNA using a partial Illumina Read 1 sequencing primer (present on every fragment in the full length cDNA library), and a reverse primer specific to exon 1 of the *Rosa26* locus.

FWD primer: 5’-CTACACGACGCTCTTCCGATCT-3’

REV primer: 5’-CTAGGTAGGGGATCGGGACT-3’

In a second round, overhanging forward primers completes the Read 1 sequencing primer binding site on the 5’ end, and biotinylated LoxCode specific primers (located inside the LoxCode construct) further enriches LoxCode cDNA.

FWD primer: 5’-AATGATACGGCGACCACCGA-GATCTACACTCTTTCCCTACACGACGCTC-3’

REV primer: 5’-biotin-TCTAGAGGATCCCCGGGTACCGA-3’

In a next round, biotinylated amplicon is purified using MyONE C1 streptavidin beads (Thermofisher), removing unwanted DNA. Then LoxCode specific DNA was cleaved off the beads using an Eco53KI restriction site 5’ of the LoxCode sequence. Finally, adaptors (IDT) were ligated using Blunt/TA ligation mix (NEB) and amplified using generic P5 and P7 primers. Libraries were sequenced on an Illumina MiSeq 600 cycle kit using custom sequencing primers.

### LoxCode Analysis

All analyses were performed in ‘R 4.1.3’ and RStudio, except for LoxCode alignment and agent-based simulations of tissue development (Fig. 4). Most graphs were created using the R package ‘ggplot 2 3.3.6’ ^55^.

### LoxCode Alignment

Raw Illumina MiSeq paired-end sequencing reads were aligned to LoxCode element sequences using custom C++ code and the library edlib^56^. Depending on the sequencing configuration, for full length LoxCodes with 13 barcode elements, 7 elements were determined from Read 1 and the remaining 6 elements from Read 2. For LoxCodes of size less than 13, a consensus barcode was determined, by comparing overlapping elements from Read 1 and Read 2. Average barcode reads for embryo 1, 2, 3, were 136K, 138K, and 130K per sample respectively. Barcode reads for all samples are provided in Supplementary Fig. 2e.

### Bulk barcode filtering, normalization and scaling

Several filtering steps were performed on the LoxCode alignment output in order to ensure only high-quality samples and barcodes were included in the analysis. Samples with less than 500 barcode reads were discarded (Supplementary Fig. 3a). Barcodes with repeated elements (likely due to PCR artifacts) and barcodes detected in only one of two PCR replicates were discarded, PCR duplicates normalized to counts per million (CPM), and barcode counts merged by taking the mean across the two samples. Next a CPM cut-off of 0.1 was applied to remove barcodes with very low frequencies. To filter out low-complexity LoxCodes for bona-fide clonal fate analysis, an empirical exclusion list of close to 30K common barcodes was generated from a total of 37 independent DNA samples from LoxCode-Rosa26CreErt2 cells (>50K cells per sample) induced at a similar recombination rate (average number of recombination: 5.14 vs 4.58 in actual data) in the embryos but in adult mice. Barcodes detected in more than 10 out of the 37 independent samples were excluded from the analysis. This cut-off provided a good compromise between stringency and the number of barcodes retained in the analysis. In some instances highlighted in the text, only barcodes detected in at least two tissues were considered for the analysis (Fig. 2e-g).

### Biomass estimation and random sampling

To evaluate the biomass for each clone (Fig. 2d), relative DNA concentrations were quantified, except for blood, from the cellular lysates using a fluorometric assay (Qubit, Invitrogen). Based on the assumption that an E12.5 embryo contains approximately 6 million cells^3^, the cell count for each clone was inferred by multiplying the barcode frequency present in the tissue sample by the relative DNA concentration and the total cell number.

In order to evaluate the presence of a clone across multiple tissues against a null hypothesis, which posits that cells are distributed randomly and uniformly across tissues according to tissue size, random samples were generated by employing the estimated cell count per clone and the relative cell count for each tissue. A comparison of the distribution of the number of tissues detected for both empirical and randomly sampled data was performed using a two-sided Kolmogorov-Smirnov test to assess the statistical significance of observed differences.

### Metaclone similarity analysis (MCSA)

Barcode counts per sample were log(1+x) transformed, and the average cosine similarity (E((A • B) / (||A|| ||B||))) between samples and across embryos (with missing samples removed) calculated and normalized to range from zero to one. Meta-clonal similarity (MCS) values were then ordered using hierarchical clustering (method=”ward.D2”) and plotted as a heatmap using the R package ‘pheatmap 1.0.12’.

### UMAP (tissues)

MCS values were computed as for the heatmap and projected onto 2 dimensions through Uniform Manifold Approximation and Projection using the R package ‘umap 0.2.7.0’ (parameters: n_neighbours_=20, spread_=2.0, min_dist_=0.1, metric=”cosine”).

### Anatomical tissue connectivity map

An anatomical 2D tissue map was created using the locations of the analysed embryonic tissues and regions. A 6-nearest-neighbour network based on MCS values was then projected onto this map connecting nodes between most similar tissues and regions. Edges with an MCS <0.75 were removed. For the purpose of visualization, edges between nodes were drawn using the edge bundling algorithm using the R package ‘ggraph 2.0.5’. To ease bundling intermediate nodes (not shown in Fig. 3g) were introduced for brain, neural tube, mesenchyme, organs, forelimbs, hindlimbs and blood.

### UMAP (barcodes)

Barcode CPM counts across tissues were log(1+x) transformed and projected onto 2 dimensions through Uniform Manifold Approximation and Projection using the R package ‘umap 0.2.7.0’ (parameters: n_neighbours=10, spread=1.0, min_dist=0.2, metric=”cosine”). Leiden clustering was performed from umap’s computed NN-network using the R package ‘leiden 0.4.3’ (resolution_parameter = 1.0).

### Comparison of bona-fide vs MCSA analysis

The Robinson-Foulds distances between trees generated from hierarchical clustering were computed using the Treedist function from the R package ‘phangorn 2.10.0’.

### Agent based modelling and parameter fitting

Tissue development was modelled using an agent-based approach written in C++ programming language, by combining previously published implementations of stochastic cellular growth^40^ and LoxCode barcoding^29^ with fate commitment. Heterogeneity in cell division times^57^, random recombination during LoxCode barcoding and fate commitment over time were assumed independent random processes, with non-synchronous divisions modelled in continuous time, barcoding occurring at a single time-point (i.e., E5.5), and for ease of implementation differentiation happening at discrete 1 hour time intervals. Times between divisions were assumed log-normal distributed (mean=12 hours, variance=2.56 hours squared), except for the first two divisions which have been shown to last approximately 20 hours^58^. For blood progenitors an approximately two-times slower division time (a factor 1.3 in the parameters of the log-normal distribution) was assumed to prevent blood becoming the dominant tissue at E12.5. Recombination between pairs of LoxP sites were chosen uniformly from the original or already recombined LoxCode cassette with LoxP sites at a distance >82 nts^29^ in the same orientations leading to inversions and LoxP sites in opposite orientations leading to excisions (number of recombination events = 4). Fate commitment was assumed to follow the hierarchy shown in Fig. 3c, with two parameters per branch point determining the minimal maturation time from the previous branchpoint (or time of fertilization for the zygote) and the probability *p* to differentiate in one or the other branch. Beyond the minimal maturation time, the chance to differentiate was assumed constant per time. In total the model has 38 parameters (32 for the differentiation process, four for division time distributions (1 for the duration of the first two divisions, 2 for the log-normal distribution, 1 for the scaling factor for blood progenitors), one for the differentiation rate, and one for the number of LoxCode recombinations). The simulations were initiated after the first two divisions (4 cells) and run until E12.5, reaching typically between 6-7 million individual agents.

To fit 33 free model parameters (parameters associated with division times and LoxCode recombination were kept fixed) to the experimental data, we utilized the covariance-matrix adaptation evolutionary strategy (CMA-ES) as outlined in reference^41^. This strategy was implemented using the C++ library libcmaes (https://github.com/CMA-ES/libcmaes). CMA-ES is a robust algorithm known for its ability to handle noise and scale effectively to high-dimensional problems. By minimizing the sum-of-squares between the entries in the empirical and simulated MCSA matrix ((17^2-17)/2-1=135 degrees of freedom) and the relative number of barcodes per tissue (16 degrees of freedom), we identified the parameters that optimally reproduced the empirical observations. For performance, in some optimizations, the MCSA matrix was evaluated 250 hours post conception rather than 300 hours. From 20 random initialization, we selected the 5 best-performing runs, all of which reached apparent convergence after 3,000 evaluations (Supplementary Fig. 5b). The best parameters were subsequently plotted (Supplementary Fig. 5c).

### Circular pedigree plots

Embryo-scale pedigree charts were produced by employing two simple rules to assign a binary code to each cell in the simulated pedigree. Firstly, the zygote received the binary code ’0’, while daughter cells inherited the same binary code as their mother, augmented by an additional digit (’0’ or ’1’). The fate at the time of division was also recorded together with the binary code. Subsequently, cells were classified by generation, arranged lexicographically, and displayed as stacked histograms with the generation on the x-axis (parameters: position=’fill’ and width scaled to generation) using circular coordinates within the ggplot2 R package and color according to fate at time of division. To generate circular pedigrees for barcoded cells at approximately E5.5, cells that underwent equal or fewer than 10 divisions (the average number of divisions at E5.5) since the zygote stage were filtered from the dataset prior to plot generation.

### Single Cell RNA sequencing analysis and cluster annotation

Sequencing data was aligned to mouse genome (mm10) with the cellranger software (version 5.0) and analysed using the R package ‘Seurat 4.0.4’. Samples were demultiplexed using the R package ‘demultiplex 1.0.2’ and custom code. Transcriptional clusters were annotated using established marker genes from the literature (Supplementary Data 2). E12.5 embryo cells could be found in two main clusters: a *Tubb2b*^+^ neuronal cluster and a mesenchymal cluster with high expression of *Col3a1, Col2a1, Col9a1, Col1a1, Prrx1, Foxd1*. Additional clusters corresponded to neural crest (*Sox10*), blood subsets (primitive erythrocytes (*Hbb-bh1, Hbb-y, Hba-a1, Hba-a2, Alas2, Gypa*), definitive erythrocytes (*Hbb-bh1-, Hbby-, Hba-a1, Hba-a2, Alas2, Gypa*), macrophages (*Cx3cr1, Csf1r, C1qa, C1qb, C1qc*), progenitors (*Mpo, Fcgr3*) and megakryocytes (*Pf4, Cd9, Itga2b, Gp1bb*), striated muscle (skeletal (*Myf5, Myod, Myog, Acat1, Tnnt1*) and cardiac (*Actc1, Tnnc1, Nppb*)), endothelium (*Cdh5, Cldn5, Kdr, Pecam1, Ramp2, Cd93*), epithelial cells (*Krt5, Krt8, Krt18, Epcam*) and pericytes (*Pdgfrb, Kcnj8, Rgs5*). Tissue of origin was consistent with transcriptional identity.

The neuronal cluster could be further subsetted into neuronal/glial progenitors (*Sox2, Nes, hes5, Btg2, Ub2c, Cenpf*), neuroblasts (*Sox2, Tubb3, Neurog2*) and differentiating neurons (*Tubb3, Stmn2, Stmn3, Sox11*). Differentiating neurons, could be broadly grouped as *Nrx3n*-(differentiating neurons 1) and *Nrx3n*+ (differentiating neurons 2) populations, with GABAergic neurons (*Dlx5*) forming a distinctive loop within the *Nrx3n*+ neurons. Neuronal neural crest derivatives (dorsal root ganglia (*Isl1, Neurod1*) and enteric neurons (*Isl1, Phoxa1, Phoxa2, Ret, Th*)) were also identifiable.

The mesenchymal cluster contained differentiating visceral and musculoskeletal cells populating the upper and lower region of that cluster, respectively (Fig. 5a). A largely undifferentiated/proliferative mesenchymal population (absent of differentiation markers, *Cenpf, Ube2c, Top2a*) was found at the interface between visceral and musculoskeletal cell types. Within the lower section of the musculoskeletal region, undifferentiated musculoskeletal mesenchyme (*Msx1, Hoxa13, Hoxd13, Lhx2*), differentiating chondrocytes (*Col2a1, Col9a1, Col9a2, Col9a3, Sox9, Acan*), tenocytes/ligamentocytes (*Scx, Dcn, Postn, Fbn2, Col5a2, Col1a1, Col3a1*), fibroblasts (*Col1a1, Col1a2, Col3a1, Cd63, Cxcl14, Shox2, Ddr2, Tnc, Vim, Pdgfra*) and meningeal fibroblasts (*Col1a1, Col1a2, Col3a1, Cd63, Vim, Zic1, Foxd1*) could be identified. The upper visceral region of the mesenchymal cluster contained ovarian mesenchyme (*Col1a1, Col1a2, Col3a1, Tcf21, Wnt4, Wt1, Kitl, Amhr2, Wnt6*), pregranulosa cells (*Upk3b, Aldh1a2, Wt1, Krt18, Krt7, Krt19, Upk1b, Alcam*), smooth muscle (*Acta2, Tagln, Myh11, Cnn1, Myocd*), pulmonary/gastrointestinal mesenchyme (*Foxf1, Wnt2, Barx1*), renal mesenchyme (*Foxd1, Pbx1, Meis2, Hoxd11, Hoxc10, Wt1*), nephron progenitors (*Meox1, Hoxd11*), valve/endocardial cushion (*Col1a1, Col3a1, Col1a2, Acta2, Tagln2, Eln, Tbx20*) and perivascular cells (*Col1a1, Col3a1, Col1a2, Cxcl12*).

### Single-cell MCSA

Similar to MCSA based on tissue level PCR barcoding data, frequencies of barcodes detected in single cells between cell types (instead of tissues) were log(1+x) transformed and used to compute scaled cosine similarity values. Only clones with at least five cells were included and to increase specificity the six most common barcodes were excluded (+9,-9,+1,+13,+1+2+3,+11+12+13). Cell types were then clustered using leiden clustering (R package ‘leiden 0.4.3’, resolution_parameter = 0.7) and projected onto 2 dimensions with the R package ‘umap 0.2.7.0’ (parameters: n_neighbors=6, spread=1, min_dist=0.01, metric=”cosine”).

### Statistical analysis

Chi-square and Kolmogorov-Smirnov tests were computed in R. If appropriate, p-values were adjusted for multiple comparisons using the Benjamini-Hochberg correction^59^. P-values below 0.05 were assumed statistically significant.

## Supporting information

Supplementary Information Guide

## Data availability

Unfiltered and filtered barcoding data, barcode exclusion list, and raw as well as processed scRNA data generated in this study have been deposited at Zenodo (10.5281/zenodo.7840234). A description of the files is provided as Supplementary Notes 1.

## Code availability

Custom C++ code for LoxCode alignment and agent-based simulations have been deposited at Zenedo (10.5281/zenodo.7840234). A description of the files is provided as Supplementary Notes 1.

## Acknowledgements

We thank the WEHI genomics and single cell core (SCORE), especially Casey Anttila for 10X library preparations and Daniel Brown for multiplexing, CRISPR mouse facility MAGEC, especially Andrew Kueh for mouse generation, and mouse facilities for animal husbandry. We thank Illumina, Inc for sequencing support. We thank Chris McGinnis and the Gartner lab (UCSF) for sharing MultiSeq reagents. We would also like to thank Ken Duffy for discussions and input during the early conceptual phase of the LoxCode technology. This work was funded by the CASS Foundation (SM/18/7776 for T.S.W.), National Health and Medical Research Council, Australia (2012196 (T.S.W), 1184736 (T.S.W. & S.H.N), 2009675 and 1145184 (S.H.N.), 1052195 (D.C.M.), 1129012 and 2011770 (S.T), the DHB Foundation (S.T) Human Frontiers Special Program (RGP0060/2012).

## Contributions

T.S.W. with S.H.N. conceived the LoxCode approach. D.C.M., T.S.W., and S.H.N. built the LoxCode construct with S.G.’s advice. T.S.W., C.B, D.C.M, P.T, S.T., S.H.N. conceived the study design. C.B. performed all embryonic dissections, with T.S.W. for tissue and single cell processing. T.S.W optimised library preparation and sequencing. T.S.W. performed all analysis and generated figures, with S.Z. and S.T. for the theoretical LoxCode recombination look-up table and empirical exclusion list respectively. C.B. provided cluster annotations, and T.S.W., C.B., S.T., P.T. and S.H.N. interpreted the data. T.S.W, D.C.M., S.T., and S.H.N. sourced funding. S.T. and S.H.N. supervised the study. All authors contributed to writing the manuscript.

## Supplementary Figures

**Supplementary Fig 1.**
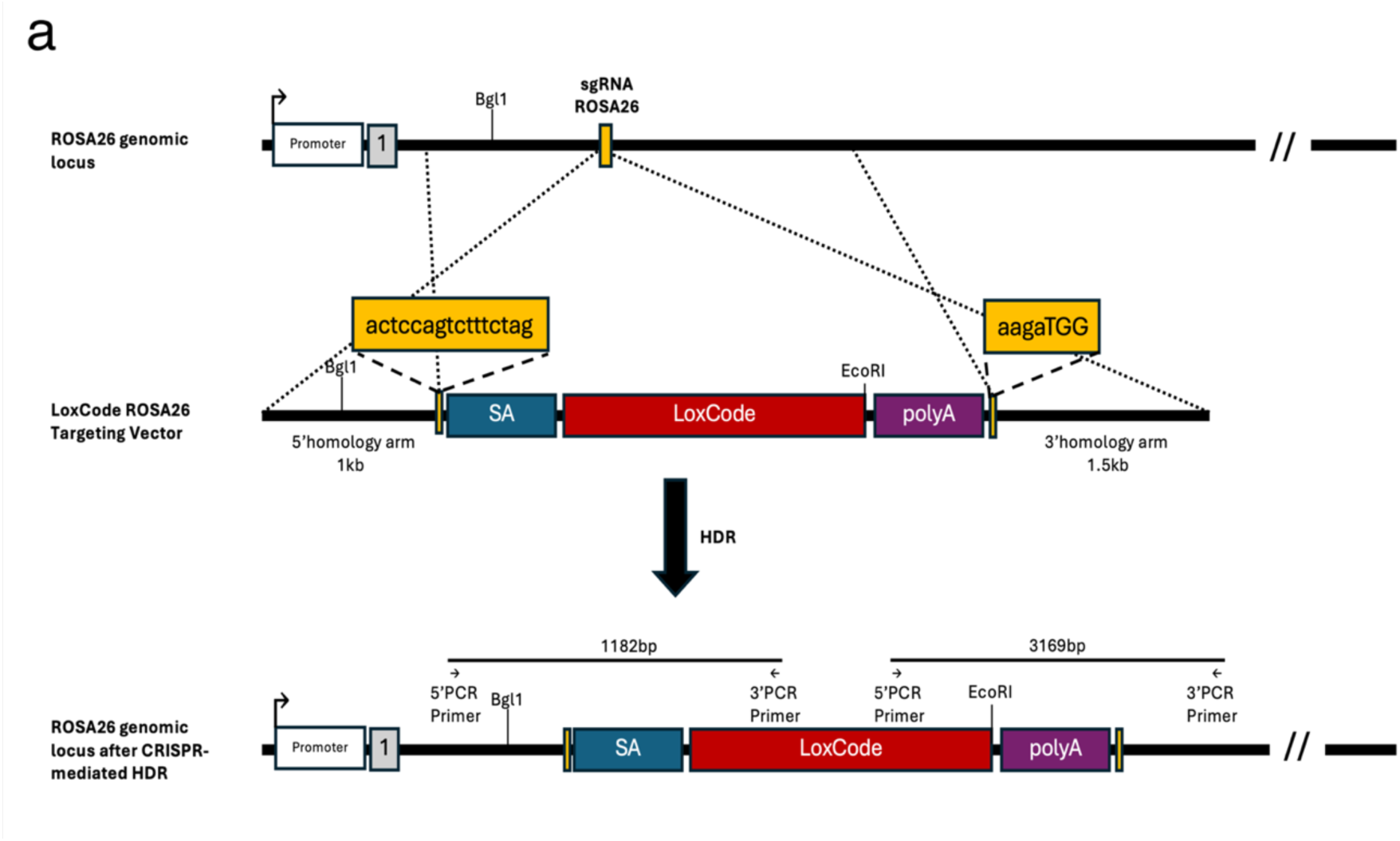
Targeting strategy for integration of the LoxCode cassette into the mouse ROS26 locus using CRISPR-Cas9 mediated homology-directed repair. Endogenous promoter, exon 1 and BglI cut site are indicated for the ROSA26 genomic locus. Integration of the LoxCode cassette occurs at the ROSA26 sgRNA binding site (yellow box). Homology-directed repair (HDR) upon CRISPR-Cas9 mediated cutting of the genome is highlighted (dotted lines) using a LoxCode ROSA26 targeting vector. The targeting vector consists of 5’ and 3’ homology arms, a splice acceptor (SA) site, the LoxCode cassette and a polyA sequence. Integration of the targeting vector disrupts the ROSA26 sgRNA binding (yellow box with sequence; dashed lines). Successful integration of the LoxCode targeting vector into the ROSA26 locus after CRISPR-mediated HDR is verified by long-range PCR followed by restriction digest of the PCR product using two sets of primer pairs, and respective restriction enzymes, BglI and EcoRI, as indicated.

**Supplementary Fig 2.**
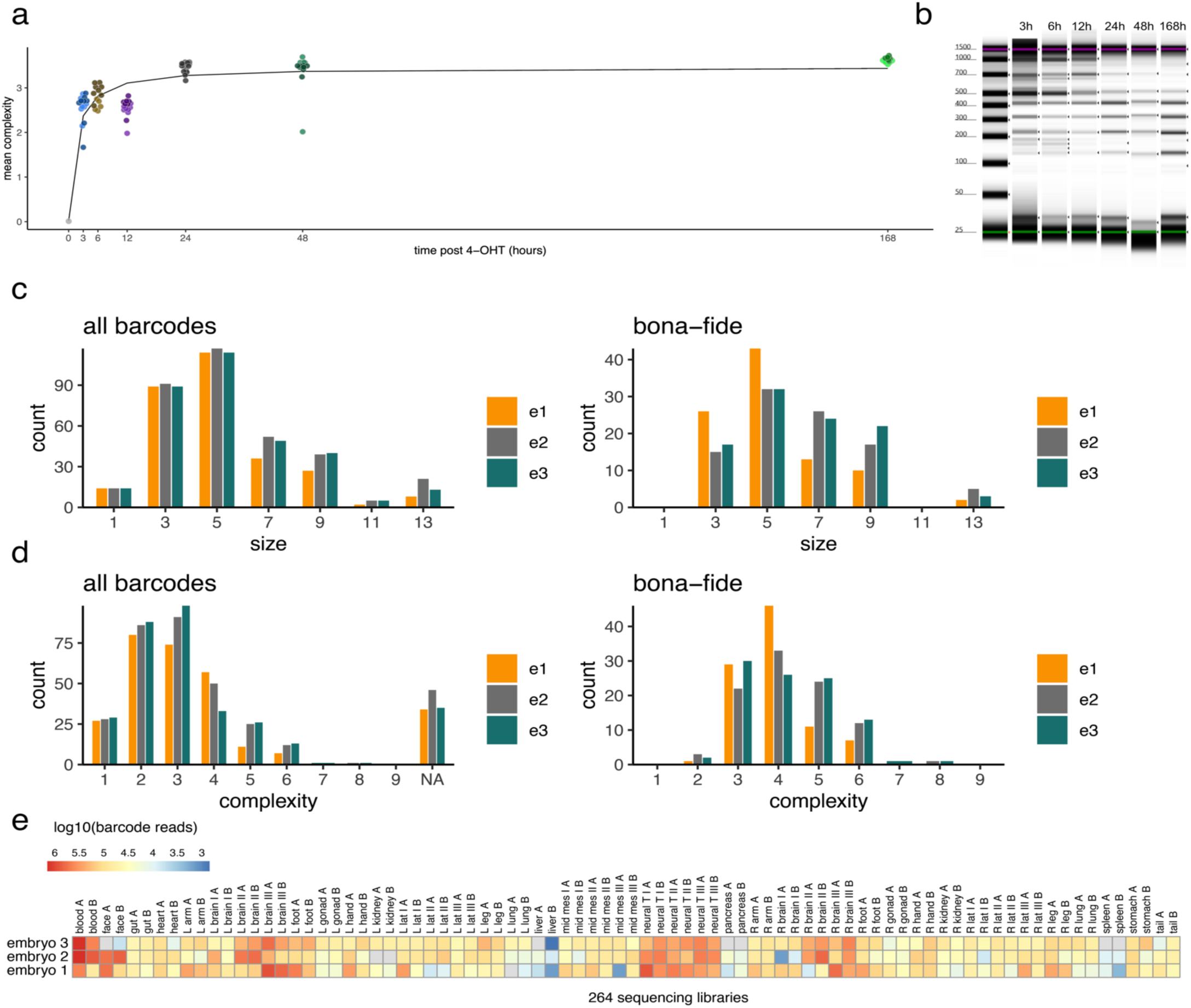
**a)** Mean Loxcode complexity 3, 6, 12, 24, 48, 168 post barcode induction in E7.5 embryos. Two litters per timepoint, each dot represents one embryo. **b)** Gel eletrophoresis traces for LoxCodes PCR amplified from embryos collected at indicated time points post tamoxifen. **c)** Distribution of barcode sizes for all (PCR-filtered) and bona-fide (empirical-frequency-filtered) clones. **d)** Distribution of the minimum number of recombinations (complexity) for all (PCR-filtered) and bona-fide clones. e) LoxCode sequencing reads for all samples (including PCR replicates).

**Supplementary Fig 3.**
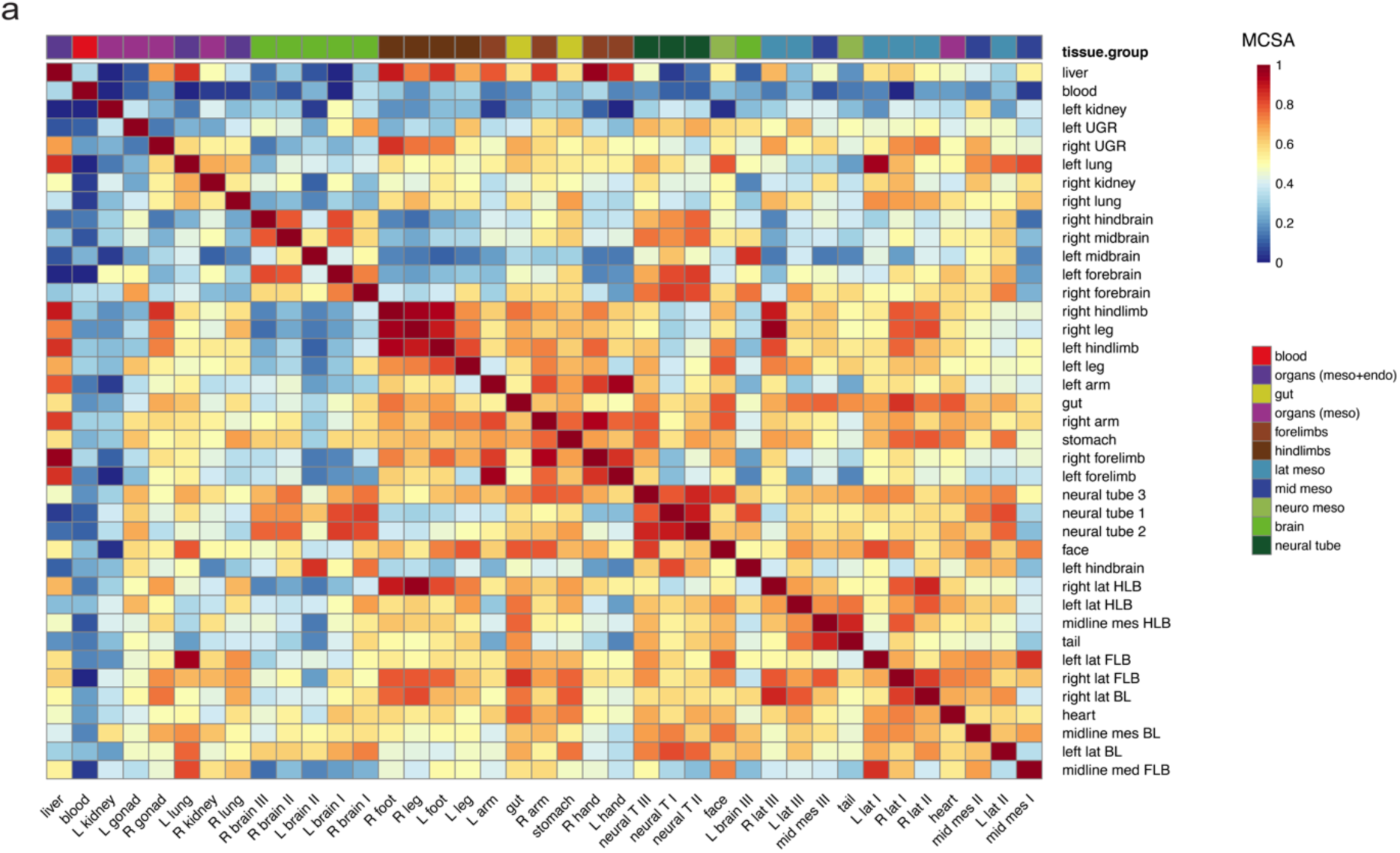
**a)** Cosine similarity matrix for all tissues using bona-fide barcodes ordered according to the corresponding MCSA heatmap in Fig. 3e.

**Supplementary Fig 4.**
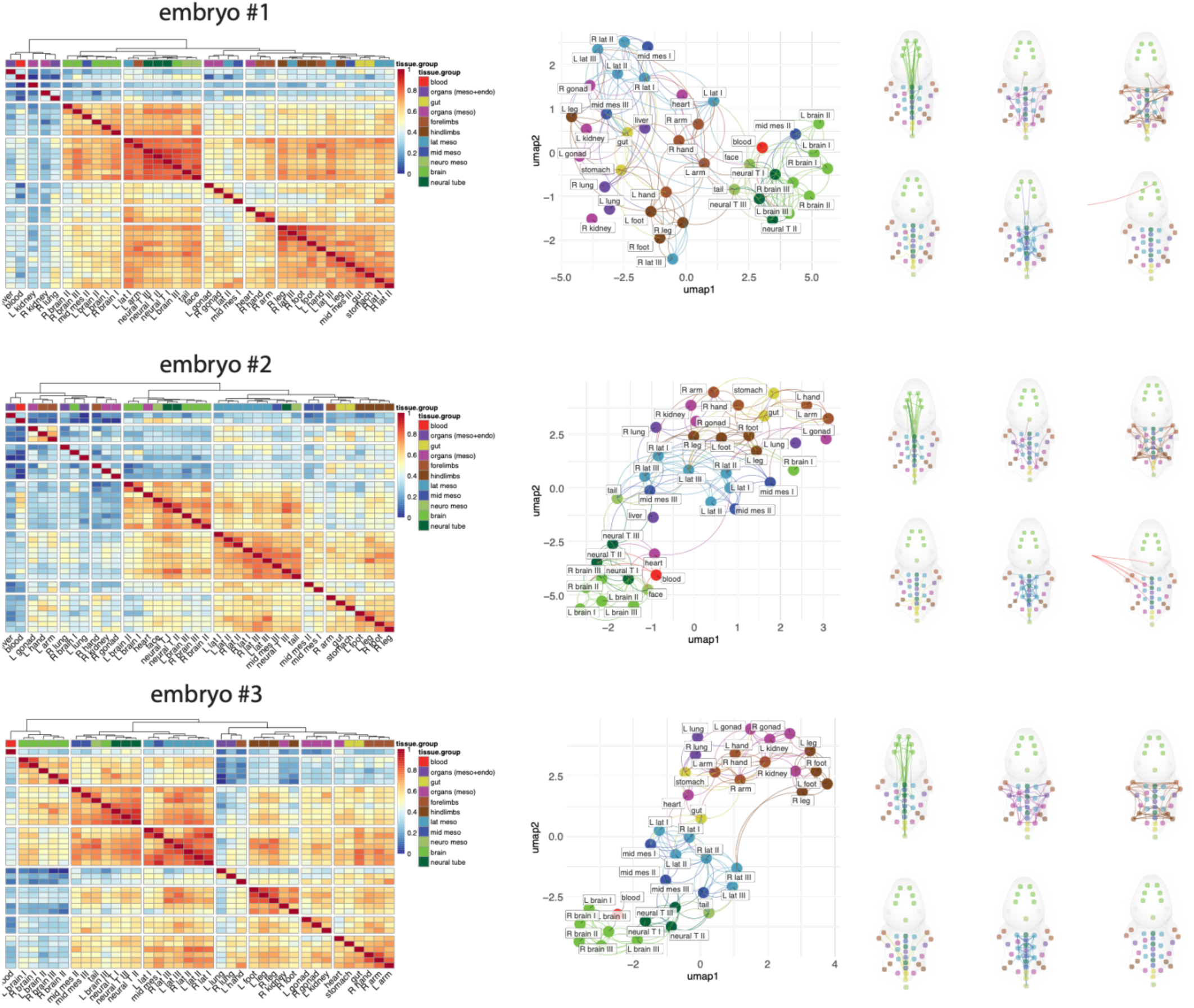
MCSA heatmap, UMAP and pseudo-anatomical map of tissue relationships. These plots are as per Figure 3 and represent the corresponding 3 biological replicates that were pooled to generate Figure 3.

**Supplementary Fig 5.**
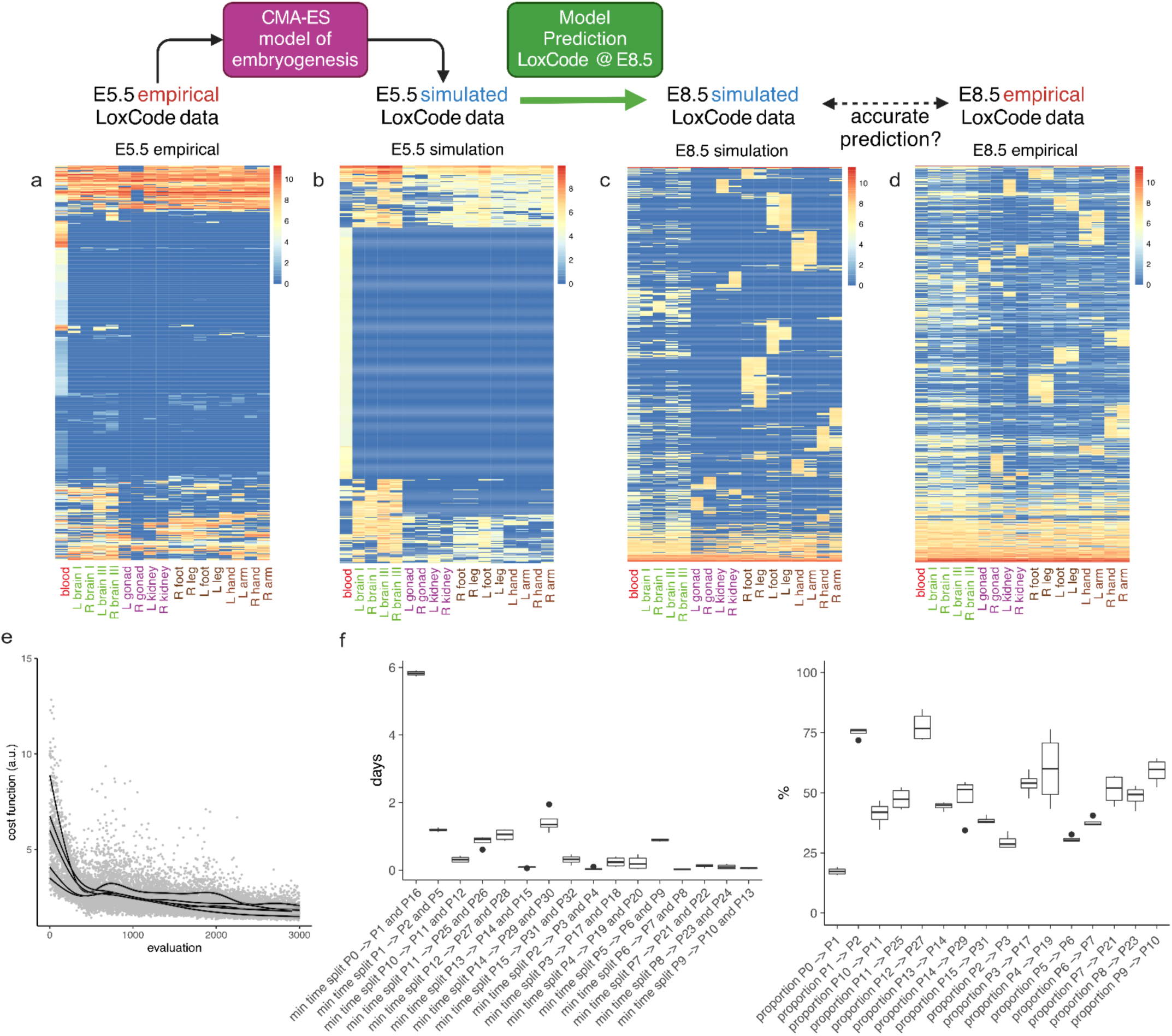
**a-d)** Heatmaps showing empirical vs simulated barcode data for LoxCode induction at E5.5 and E8.8 and embryo harvesting at E12.5. E8.5 data (empirical and simulated) is filtered for relative clone size (>0.1%) and detection in at least two tissues. **e)** Convergence of the CMA-ES optimization routine (5 out of 20 best performing runs are shown). **f)** Best-fit parameters from 5 best performing runs (boxplot indicating median, 25^th^ and 75^th^ quantile). Populations and parameters are provided in Supplementary Methods 1.

**Supplementary Fig 6.**
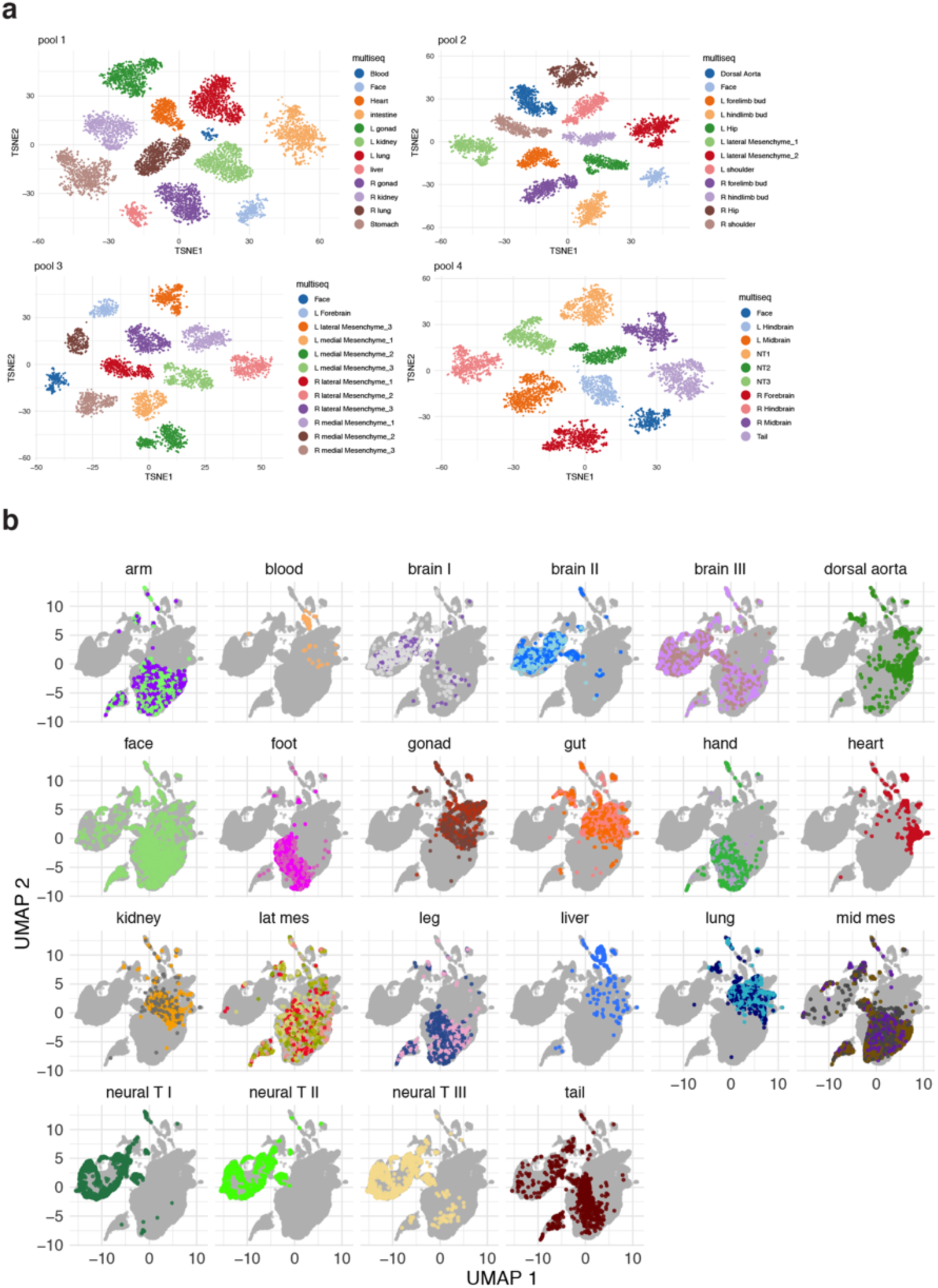
The 40 tissues, which were dissected and processed into a single-cell suspension as depicted in Figure 1, underwent multiplexing using the MULTI-seq method. Subsequently, the samples were pooled and analyzed using a droplet-based scRNA-seq platform with a 3’ capture scRNA-seq kit. Demultiplexing via MULTI-seq proved effective in: a) distinguishing the tissues of origin, and b) differentially distributing them across the transcriptional space (as illustrated by the UMAP in Figure 4). It should be noted that distinct colors represent left and right samples (or I, II, III for mesenchymal tissues) of the specified tissue, such as left and right kidneys.

**Supplementary Fig 7.**
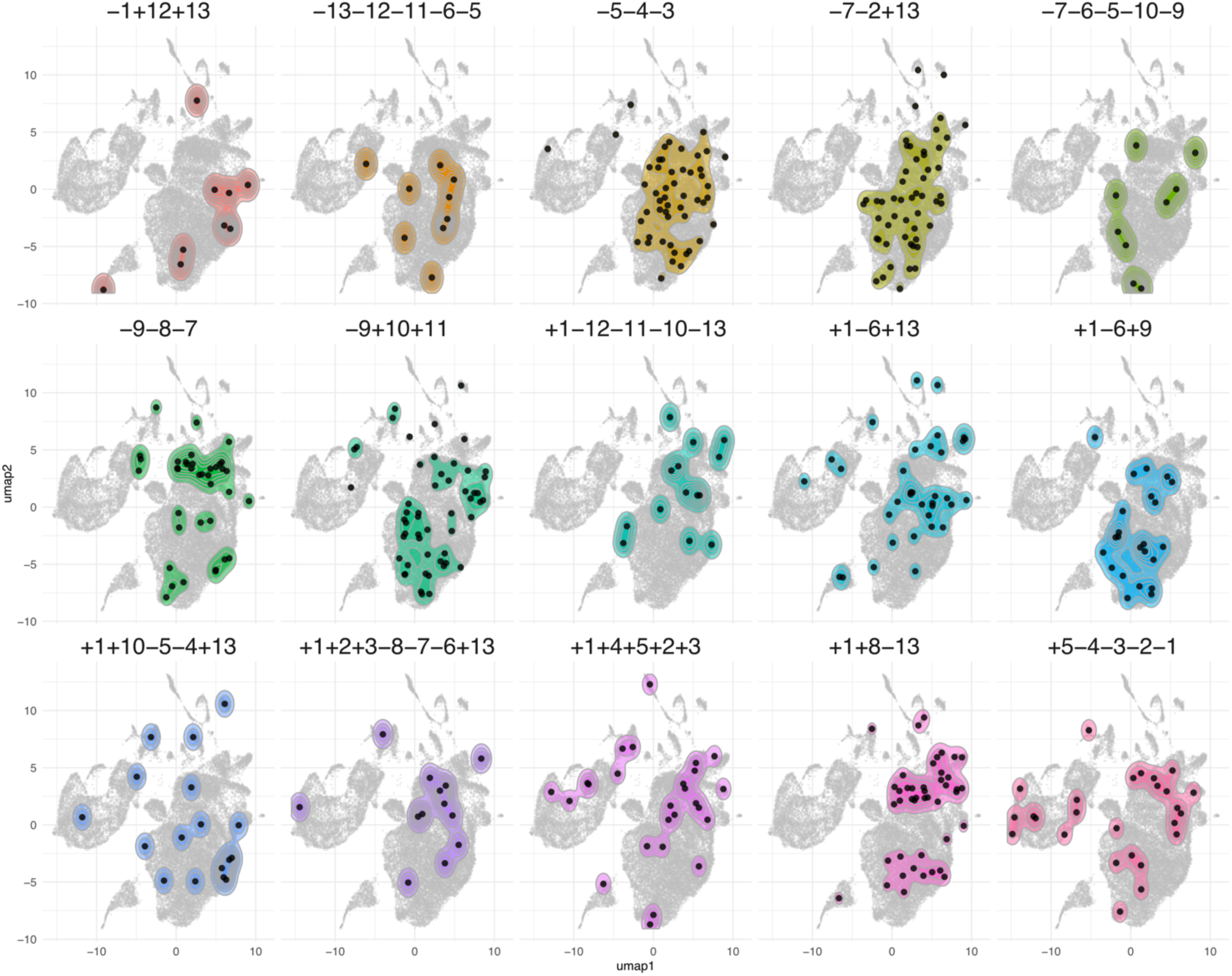
Clonal fate heterogeneity. **a)** Single cell output corresponding to the indicated LoxCodes of bona-fide clones for which more than 8 cells were captured.

**Supplementary Fig 8.**
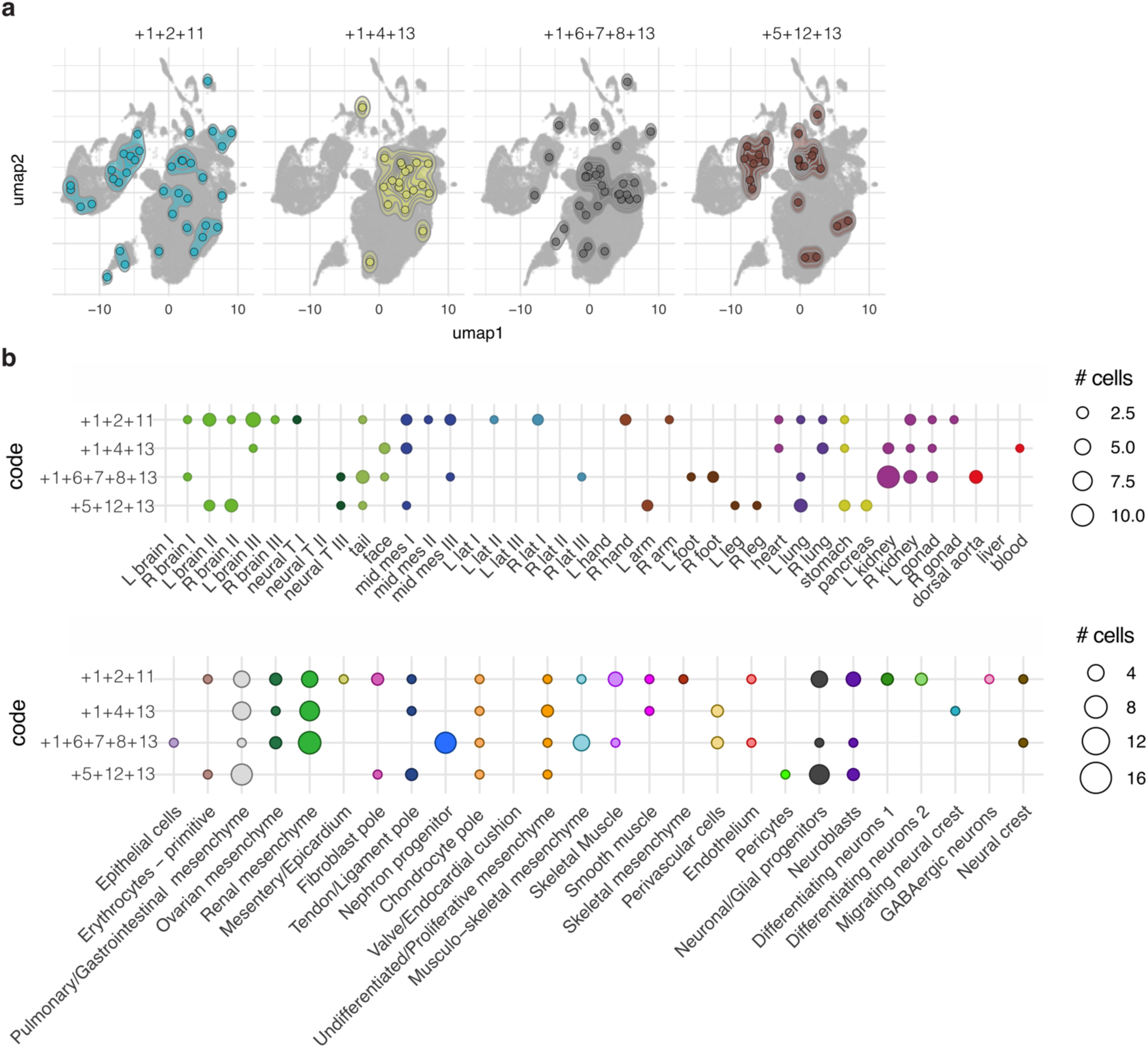
Meta-clonal fate heterogeneity. **a)** Examples of single cell meta-clonal output corresponding to the indicated LoxCodes. **b)** Distribution of single cells per detected LoxCode according to **t**issue (top) or cell type (bottom).

**Supplementary Fig 9.**
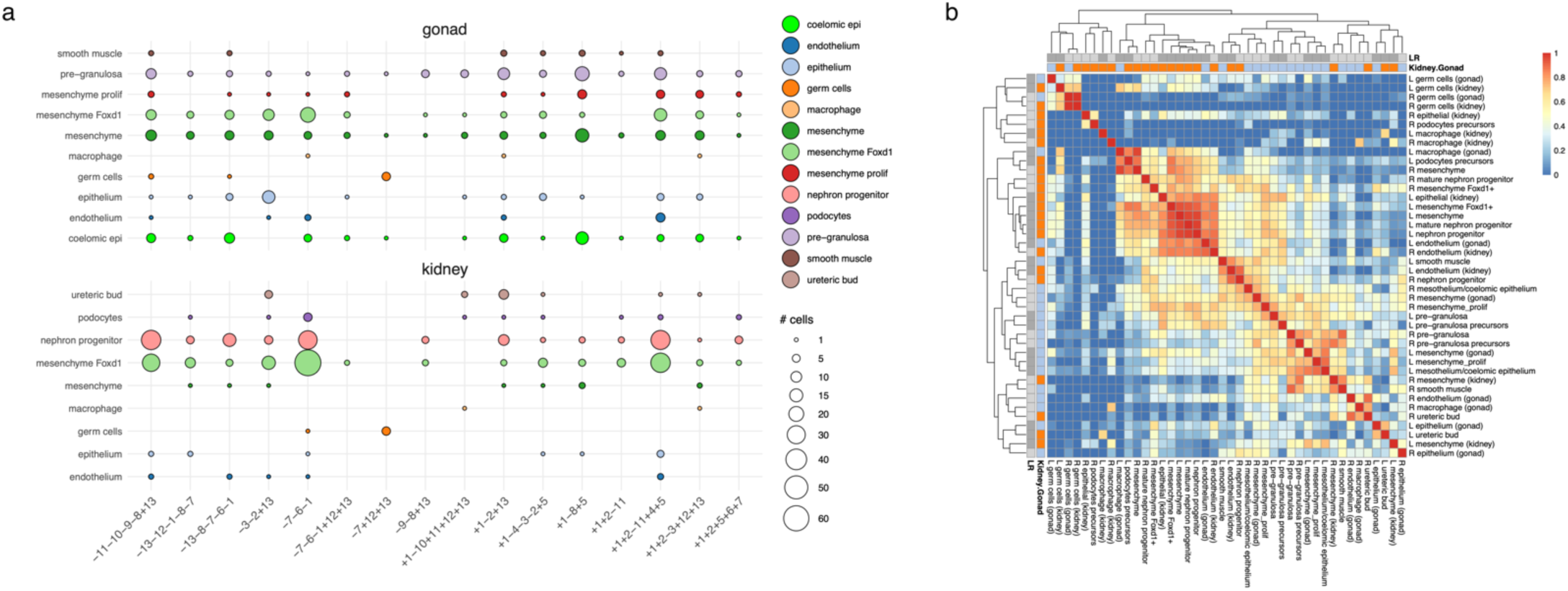
Heterogeneity of epiblast contribution to gonads and kidneys. **a)** Clonal gonad-kidney cell-type bias (* p-value<0.05, ** adj p-value <0.05). Colour indicates whether the cell type is over- or underrepresented, the dot size is proportional to the number of cells captured. **b**) scCSA between transcriptional clusters in left and right gonad and kidney.

