## Supplementary Information Guide for "LoxCode in vivo barcoding resolves epiblast clonal fate to fetal organs"

Supplementary Methods 1

Description of model populations (from P0 to P32) and parameters (p1 to p38). For variable parameters (p1 to p33), mean and standard deviation of five best performing optimization runs are provided. Parameters p1 to p16 and p33 are in units of hours, p17 to p32 are unitless.

Supplementary Notes 1

Description of data and C++ code deposited at Zenodo (10.5281/zenodo.7840234).

Supplementary Data 1

Scalable vector graphics of circular pedigree plot showing *in silico* embryogenesis from zygote to E12.5.

Supplementary Data 2

Details on transcriptional cluster annotation of single-cell data in Fig. 5a (sheet 1: DE genes per cluster, sheet 2: cluster markers and references).
