## Supplementary Information for "LoxCode in vivo barcoding resolves epiblast clonal fate to fetal organs"

### Supplementary Methods 1

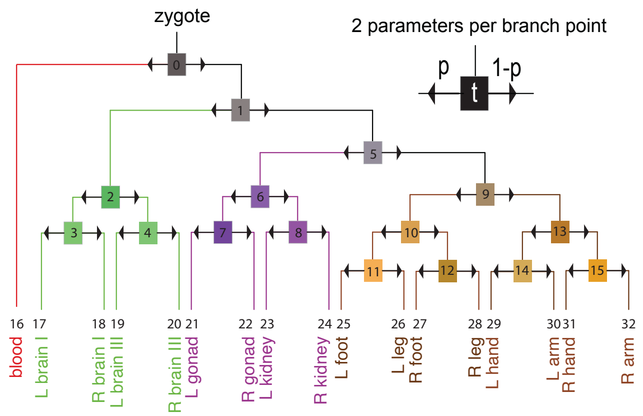

presumptive progenitors (model)

- 0 Epiblast (pre-gastrulation)
- 1 Epiblast (early gastrulation)
- 5 Mesendoderm progenitor
- 2 Neural progenitor
- 3 CNS forebrain neural progenitor
- 4 CNS hindbrain neural progenitor
- 6 Intermediate mesoderm progenitor
- 7 Mesonephric progenitor
- 8 Metanephric progenitor
- 9 Lateral mesoderm progenitor
- 10 Hindlimb progenitor
- 13 Forelimb progenitor
- 11 Hindlimb progenitor (left)
- 12 Hindlimb progenitor (right)
- 14 Forelimb progenitor (left)
- 15 Forelimb progenitor (right)

| cell type | population (model) |
|---|---|
| Epiblast (pre-gastrulation) | P0 |
| Epiblast (early gastrulation) | P1 |
| Mesendoderm progenitor | P5 |
| Neural progenitor | P2 |
| CNS forebrain neural progenitor | P3 |
| CNS hindbrain neural progenitor | P4 |
| Intermediate mesoderm progenitor | P6 |
| Mesonephric progenitor | P7 |
| Metanephric progenitor | P8 |
| Lateral mesoderm progenitor | P9 |
| Hindlimb progenitor | P10 |
| Forelimb progenitor | P13 |
| Hindlimb progenitor (left) | P11 |
| Hindlimb progenitor (right) | P12 |
| Forelimb progenitor (left) | P14 |
| Forelimb progenitor (right) | P15 |
| blood | P16 |
| L brain I | P17 |
| R brain I | P18 |
| L brain III | P19 |
| R brain III | P20 |
| L gonad | P21 |
| R gonad | P22 |
| L kidney | P23 |
| R kidney | P24 |
| L foot | P25 |
| L leg | P26 |
| R foot | P27 |
| R leg | P28 |
| L hand | P29 |
| L arm | P30 |
| R hand | P31 |
| R arm | P32 |

| variable parameters: | | mean | sd |
|---|---|---|---|
| p1 | min time split P0 -> P1 and P16 | 139.9 | 1.9 |
| p2 | min time split P1 -> P2 and P5 | 28.6 | 1.4 |
| p3 | min time split P2 -> P3 and P4 | 1.1 | 0.9 |
| p4 | min time split P3 -> P17 and P18 | 5.8 | 3.7 |
| p5 | min time split P4 -> P19 and P20 | 5.3 | 5.1 |
| p6 | min time split P5 -> P6 and P9 | 21.4 | 1.0 |
| p7 | min time split P6 -> P7 and P8 | 0.5 | 0.4 |
| p8 | min time split P7 -> P21 and P22 | 2.9 | 1.3 |
| p9 | min time split P8 -> P23 and P24 | 2.4 | 1.7 |
| p10 | min time split P9 -> P10 and P13 | 1.4 | 0.6 |
| p11 | min time split P10 -> P11 and P12 | 7.5 | 2.2 |
| p12 | min time split P11 -> P25 and P26 | 20.5 | 4.0 |
| p13 | min time split P12 -> P27 and P28 | 25.1 | 4.2 |
| p14 | min time split P13 -> P14 and P15 | 2.1 | 0.4 |
| p15 | min time split P14 -> P29 and P30 | 34.5 | 8.6 |
| p16 | min time split P15 -> P31 and P32 | 7.5 | 3.4 |
| p17 | proportion differentiate P0 -> P1 | 17.3 | 1.5 |
| p18 | proportion differentiate P1 -> P2 | 75.1 | 2.2 |
| p19 | proportion differentiate P2 -> P3 | 29.6 | 3.2 |
| p20 | proportion differentiate P3 -> P17 | 53.8 | 4.9 |
| p21 | proportion differentiate P4 -> P19 | 59.9 | 15.3 |
| p22 | proportion differentiate P5 -> P6 | 30.7 | 1.4 |
| p23 | proportion differentiate P6 -> P7 | 37.7 | 1.9 |
| p24 | proportion differentiate P7 -> P21 | 51.3 | 6.4 |
| p25 | proportion differentiate P8 -> P23 | 48.5 | 4.6 |
| p26 | proportion differentiate P9 -> P10 | 59.0 | 5.4 |
| p27 | proportion differentiate P10 -> P11 | 41.3 | 5.1 |
| p28 | proportion differentiate P11 -> P25 | 47.5 | 4.5 |
| p29 | proportion differentiate P12 -> P27 | 77.6 | 6.3 |
| p30 | proportion differentiate P13 -> P14 | 44.4 | 1.7 |
| p31 | proportion differentiate P14 -> P29 | 47.9 | 9.2 |
| p32 | proportion differentiate P15 -> P31 | 38.5 | 1.6 |
| p33 | differentiation rate | 5.5 | 0.7 |
| fixed parameters: | | | |
| p34 = 20 hours | division time of first two divisions | | |
| p35 = 12 hours | average division time (all except blood progenitors) | | |
| p36 = 2.54 | std division time (all except blood progenitors) | | |
| p37 = 1.3 | factor for blood progenitor division time | | |
| p38 = 4 | number of LoxCode recombinations | | |

### Supplementary Notes 1

#### Data and C++ code deposited at Zenodo (10.5281/zenodo.7840234)

##### bulk

Bulk analysis data

- barcoding_data
- exclusion_list
- loxcode_alignement

##### modelling

Modelling data

- C++
- pedigree

##### single_cell

Single cell analysis data

- processed
- raw

#### Descriptions

##### barcoding_data

Barcoding data files

- barcodes_post_exclusion_filtering_two_tissues.xlsx: Barcodes detected in at least two tissues
- barcodes_post_exclusion_filtering.xlsx: Barcodes after exclusion filtering
- barcodes_post_PCR_filtering.xlsx: Barcodes after PCR filtering
- barcodes_post_PCR_merging.xlsx: Barcodes after PCR merging
- barcodes_pre_filtering.xlsx: Barcodes before any filtering

##### exclusion_list

Exclusion list data

- exclusion_list.xlsx: Exclusion list in Excel format. First column indicates barcode, the second column the number of samples (from a total of 37 reference samples) the barcode is detected in.

##### loxcode_alignement

Loxcode alignment data

- edlib: Edlib library for sequence alignment
  - include: Header files for the library
    - edlib.h: Main header file for Edlib

    - src: Source files for the library
        - edlib.cpp: Main source file for Edlib
- loxcode_align.cpp: Source code for loxcode alignment program
- output: Output files from loxcode alignment
    - sample_1_loxcode_counts.csv: Loxcode counts for sample 1
    - sample_2_loxcode_counts.csv: Loxcode counts for sample 2
- test_fastq: example FASTQ files
    - sample_1_R1_001.fastq: Read 1 of sample 1
    - sample_1_R2_001.fastq: Read 2 of sample 1
    - sample_2_R1_001.fastq: Read 1 of sample 2
    - sample_2_R2_001.fastq: Read 2 of sample 2

### C++

C++ source code and libraries

- loxcode_simulate_embryogenesis.cpp: Source code for loxcode embryogenesis simulation
- model_populations_parameters.xlsx: Model populations (sheet 1) and parameters (sheet 2) in Excel format

##### pedigree

Pedigree data

- loxcode_census_at_E12_5.csv: simulated Loxcode census at E12.5
- loxcode_census_at_E5_5.csv: simulated Loxcode census at E5.5
- loxcode_full_pedigree.csv: Full pedigree

##### processed

Processed single cell data

- seurat: Seurat analysis data
    - seurat_object.rds: Seurat R object with meta data including single-cell LoxCode information and tissue of origin
- web_summaries: Web summaries of single cell data
    - web_summary_pool_1.html: 10X Genomics Web summary for pool 1
    - web_summary_pool_2.html: 10X Genomics Web summary for pool 2
    - web_summary_pool_3.html: 10X Genomics Web summary for pool 3
    - web_summary_pool_4.html: 10X Genomics Web summary for pool 4

##### raw

Raw single cell data

- fastq: FASTQ files for single cell data
    - pool_1_S1_R1_001.fastq.gz: Read 1 fastq file for pool 1
    - pool_1_S1_R2_001.fastq.gz: Read 2 fastq file for pool 1
    - pool_2_S2_R1_001.fastq.gz: Read 1 fastq file for pool 2
    - pool_2_S2_R2_001.fastq.gz: Read 2 fastq file for pool 2

    - pool_3_S3_R1_001.fastq.gz: Read 1 fastq file for pool 3
    - pool_3_S3_R2_001.fastq.gz: Read 2 fastq file for pool 3
    - pool_4_S4_R1_001.fastq.gz: Read 1 fastq file for pool 4
    - pool_4_S4_R2_001.fastq.gz: Read 2 fastq file for pool 4
- raw_feature_bc_matrix: Raw feature barcode matrices
    - pool_1: Raw feature barcode matrix for pool 1
        - barcodes.tsv.gz: Barcodes for pool 1
        - features.tsv.gz: Features for pool 1
        - matrix.mtx.gz: Matrix for pool 1
    - pool_2: Raw feature barcode matrix for pool 2
        - barcodes.tsv.gz: Barcodes for pool 2
        - features.tsv.gz: Features for pool 2
        - matrix.mtx.gz: Matrix for pool 2
    - pool_3: Raw feature barcode matrix for pool 3
        - barcodes.tsv.gz: Barcodes for pool 3
        - features.tsv.gz: Features for pool 3
        - matrix.mtx.gz: Matrix for pool 3
    - pool_4: Raw feature barcode matrix for pool 4
        - barcodes.tsv.gz: Barcodes for pool 4
        - features.tsv.gz: Features for pool 4
        - matrix.mtx.gz: Matrix for pool 4
