## Supplementary figures and images for "LoxCode in vivo barcoding resolves epiblast clonal fate to fetal organs"

### Supplementary Data 1

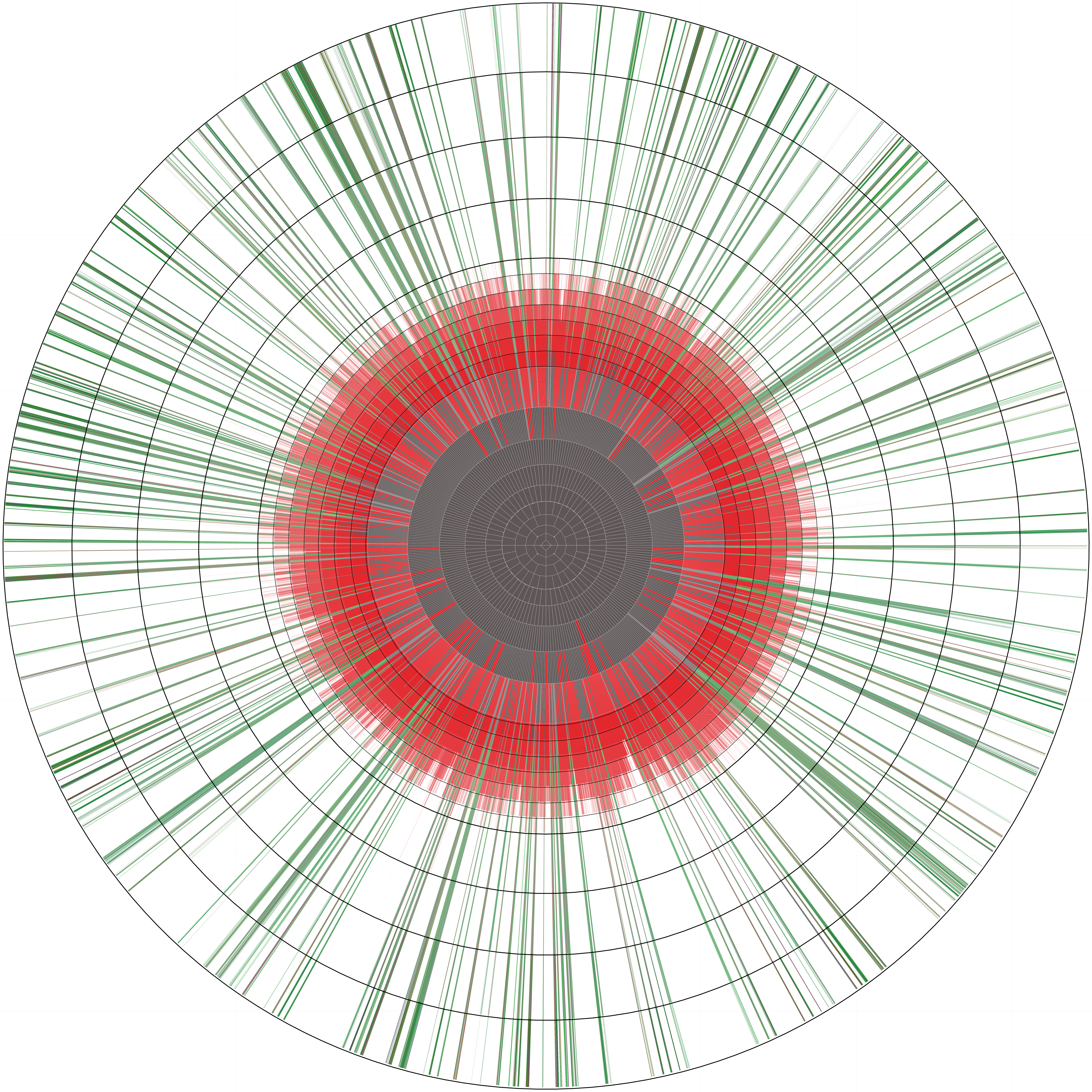
